# Evolutionary analysis of PINK1 reveals key roles of the N- and C-terminal extensions and TOM20 in folding its kinase domain

**DOI:** 10.64898/2026.09.05.749644

**Authors:** Tara Shomali, Simon Veyron, Jean-François Trempe

## Abstract

PTEN-induced kinase 1 (PINK1) is a mitochondrial serine/threonine kinase that initiates ubiquitin-dependent mitophagy and is mutated in early-onset Parkinson’s disease. Despite extensive characterization of insect PINK1 orthologues, obtaining soluble and catalytically active recombinant *Homo sapiens* PINK1 (HsPINK1) has remained challenging, limiting its biochemical and structural investigation. Here, we identify an evolutionary divergence in the mechanisms supporting PINK1 stability and activity. Comparative analysis of 48 metazoan PINK1 orthologues revealed that predicted binding to the mitochondrial import receptor TOM20 is predominantly a vertebrate feature and coincides with divergence of the PINK1 N- and C-terminal extensions (NTE and CTE). Consistent with this distinction, invertebrate PINK1 orthologues were purified as active kinases, whereas vertebrate orthologues lacked detectable ubiquitin kinase activity. NMR spectroscopy demonstrated that humanization of ten residues within the *Tribolium castaneum* (Tc)PINK1 NTE–CTE interface is sufficient to confer direct binding to human TOM20, while co-expression with TOM20 increased the activity of recombinant HsPINK1. Guided by these evolutionary differences, we engineered a chimeric PINK1 containing a predominantly human kinase domain supported by Tc-derived NTE–CTE elements. The resulting kinase was catalytically active and required compatible interactions between the NTE–CTE and kinase C-lobe for activity. The chimera enabled characterization of PINK1 variants and revealed R152W as a likely pathogenic variant. Together, our findings support a model in which the NTE and CTE act as intramolecular determinants of PINK1 kinase stability and suggest that vertebrate PINK1 has evolved an increased dependence on the mitochondrial import machinery to maintain a functional state.

## Introduction

The serine/threonine kinase PTEN-induced kinase 1 (PINK1) is a central regulator of mitochondrial quality control and an initiator of ubiquitin-dependent mitophagy (Bayne & Trempe, 2026). Loss-of-function mutations in PINK1 cause autosomal recessive early-onset Parkinson’s disease (EOPD) (Valente et al., 2004), highlighting the importance of PINK1-mediated signaling in neuronal survival. Under basal conditions, PINK1 is constitutively imported into mitochondria and rapidly degraded (Geisler et al., 2010; Matsuda et al., 2010; Narendra et al., 2010). Upon mitochondrial depolarization, import is stalled, allowing PINK1 to accumulate on the translocase of the outer mitochondrial membrane (TOM) complex (Lazarou et al., 2012), where it becomes activated through trans-autophosphorylation (Okatsu et al., 2012; Okatsu et al., 2013; Rasool et al., 2018). Activated PINK1 phosphorylates ubiquitin at Ser65, generating a receptor for the E3 ubiquitin ligase Parkin (Kane et al., 2014; Kondapalli et al., 2012; Koyano & Matsuda, 2015; Koyano et al., 2014). Subsequent phosphorylation of the Parkin ubiquitin-like (Ubl) domain activates ubiquitin ligase activity, resulting in a feed-forward amplification loop that labels damaged mitochondria for mitophagy and mitochondrial-derived vesicle (MDV) formation (McLelland et al., 2014).

Structurally, PINK1 consists of an N-terminal mitochondrial targeting sequence (MTS), a transmembrane helix contiguous with an N-terminal extension (NTE), a conserved kinase domain containing three unique insertions, and a C-terminal extension (CTE). X-ray crystallography and cryo-electron microscopy (cryo-EM) structures of insect PINK1 (Gan et al., 2022; Rasool et al., 2022) and human PINK1 bound to the TOM complex (Callegari et al., 2025) show that the NTE and CTE bind to the kinase domain and are an integral part of the cytosolic domain. Yet, despite nearly two decades of research, obtaining soluble, catalytically active recombinant human PINK1 (HsPINK1) has remained a major challenge. Consequently, most structural, biochemical, and inhibitor discovery studies have relied on insect orthologues such as *Tribolium castaneum* PINK1 (TcPINK1) and *Pediculus humanus* PINK1 (PhPINK1), which are readily expressed in bacteria and exhibit robust kinase activity (Woodroof et al., 2011). Although these orthologues have provided important mechanistic insight, they do not fully recapitulate the biochemical and regulatory properties of HsPINK1. For example, TcPINK1 exhibits different affinities towards kinase inhibitors compared to HsPINK1, tested in reconstitution assays (Rasool et al., 2024).

Recent biochemical studies identified TOM20 as a critical regulator of HsPINK1, demonstrating that it enhances kinase stability and activity in cells (Eldeeb et al., 2024). This effect is mediated by an interaction between TOM20 and the NTE-CTE region of PINK1, which was predicted by AlphaFold and tested by mutagenesis. More recently, the cryoEM structure of HsPINK1 bound to the TOM complex confirmed that TOM20 interacts directly with the PINK1 NTE-CTE regions (Callegari et al., 2025). However, it remains unclear whether this interaction is a conserved feature of PINK1 biology, and whether TOM20 binding is required for the folding and activation of PINK1.

Here, we demonstrate that the PINK1-TOM20 interaction is a vertebrate-specific evolutionary feature that fundamentally distinguishes vertebrate and invertebrate PINK1 proteins. Comparative sequence analysis and AlphaFold3 predictions identify extensive divergence within the NTE and CTE regions that coincides with the evolution of TOM20 binding. Consistent with these predictions, vertebrate PINK1 orthologues fail to express as active recombinant kinases, whereas invertebrate orthologues readily produce catalytically active protein. Using NMR spectroscopy and recombinant co-expression, we show that TOM20 binds HsPINK1 via the NTE-CTE and promotes formation of the active state of the kinase. Based on these results, we engineered an active chimeric enzyme with a predominantly HsPINK1 kinase domain fused to the NTE-CTE region of TcPINK1, which is dependent on the restoration of compatibility between the NTE-CTE module and the kinase C-lobe. Finally, we demonstrate that this engineered chimeric enzyme reproduces the biochemical consequences of EOPD mutations, providing new mechanistic insight into PINK1 activation while establishing a system for structural biology and therapeutic discovery.

## Results

### PINK1 is predicted to bind TOM20 only in vertebrates

Several studies have reported that recombinant human PINK1 is considerably less active and stable than its insect orthologues (Rasool et al., 2018; Woodroof et al., 2011). To investigate whether evolutionary divergence may underlie these differences, we compared the sequences of representative vertebrate and invertebrate PINK1 proteins (Fig. 1a). While the kinase domain proper is highly conserved, we observed considerably lower sequence conservation in the NTE-CTE regions, as well as the insert-1 and insert-2. Insert-3, which is responsible for binding ubiquitin (Kumar et al., 2017; Schubert et al., 2017), is also highly conserved. Mapping of the sequence conservation on the structures of *Homo sapiens* PINK1 (HsPINK1) and *Tribolium castaneum* PINK1 (TcPINK1) highlighted this divergence, the active site showing the most conservation, and with considerably lower conservation on the NTE-CTE side (Fig. 1b). The NTE and CTE indeed share only 14% and 11% sequence identity between the two species, respectively, whereas the kinase domain retained 46% identity (Table S1). Since the NTE–CTE region was recently identified as the TOM20-binding interface (Eldeeb et al., 2024), we hypothesized that TOM20 recognition differs across species and may have evolved specifically in vertebrates.

**Figure 1.**
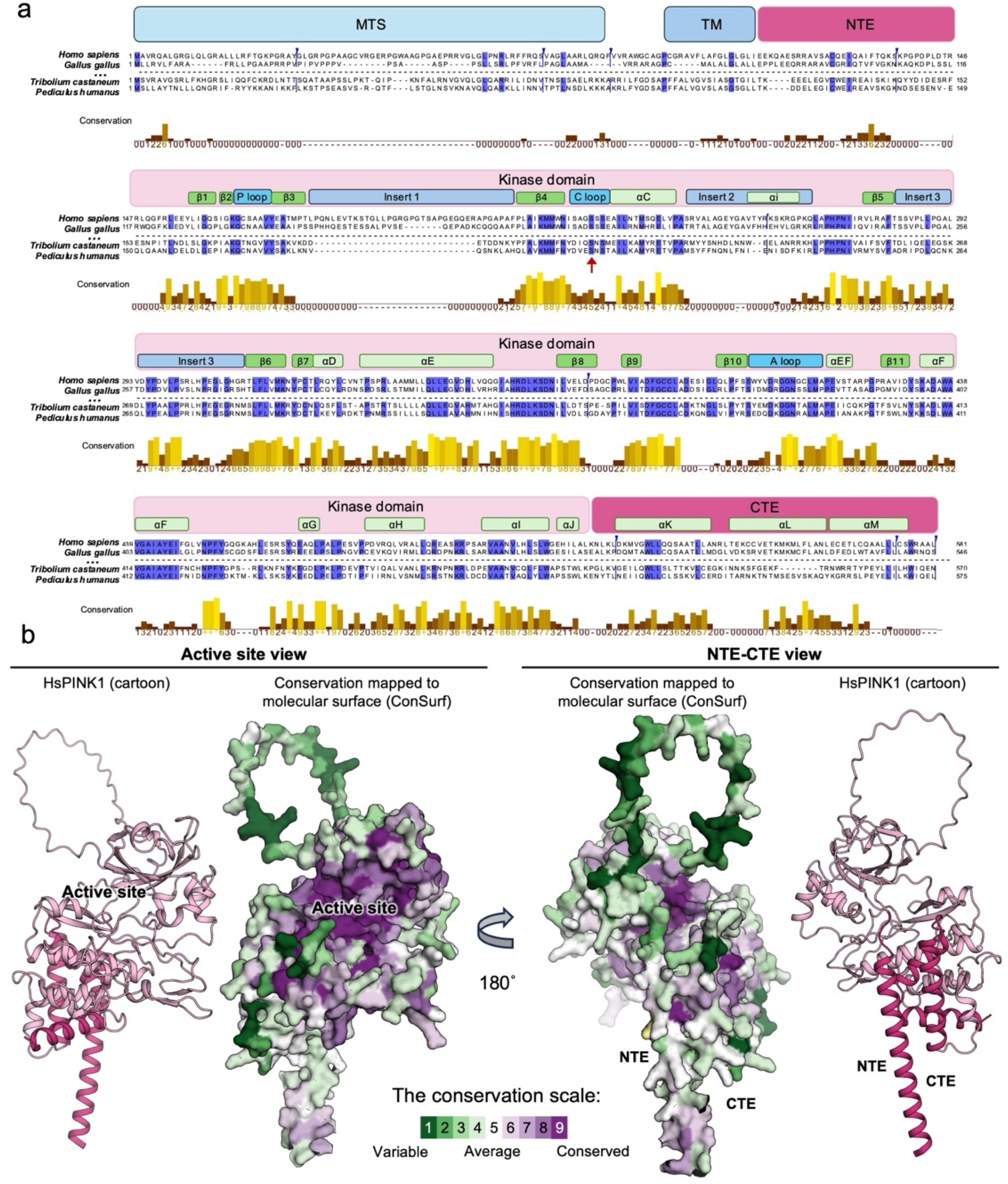
Evolutionary conservation of PINK1 domains. **(a)** Multiple sequence alignment of PINK1 from *Homo sapiens*, *Gallus gallus*, *Tribolium castaneum*, and *Pediculus humanus*, shown relative to the human PINK1 (HsPINK1) sequence. The conservation profile above the alignment represents residue conservation calculated from a multiple sequence alignment of PINK1 orthologues from 48 species spanning diverse metazoan phyla. Insertions and gaps relative to HsPINK1 were omitted for visualization. PINK1 structural regions are indicated above the sequence, and residue identity relative to HsPINK1 is highlighted. **(b)** Evolutionary conservation mapped onto the structure of HsPINK1 using ConSurf, based on the multiple sequence alignment of 48 PINK1 orthologues. Residues are coloured according to ConSurf conservation scores from variable (1) to highly conserved (9). Cartoon and molecular-surface representations are shown from the active-site and NTE–CTE-facing orientations. The N-terminal region was omitted for visualization.

To test this hypothesis, we predicted PINK1–TOM20 complexes using AlphaFold3 across a panel of 48 representative species (Fig. S1). The panel spanned major metazoan phyla, including mammals, birds, reptiles, amphibians, fish, insects, nematodes and cnidarians (Table S1). Vertebrate PINK1 orthologues consistently produced high-confidence and convergent predictions for the PINK1–TOM20 complex, whereas invertebrate orthologues, including TcPINK1, exhibited substantially lower Multimer scores, with most scoring below 0.5 (Fig. 2a, 2b). These results suggest that the ability of PINK1 to interact with TOM20 was acquired during vertebrate evolution.

**Figure 2.**
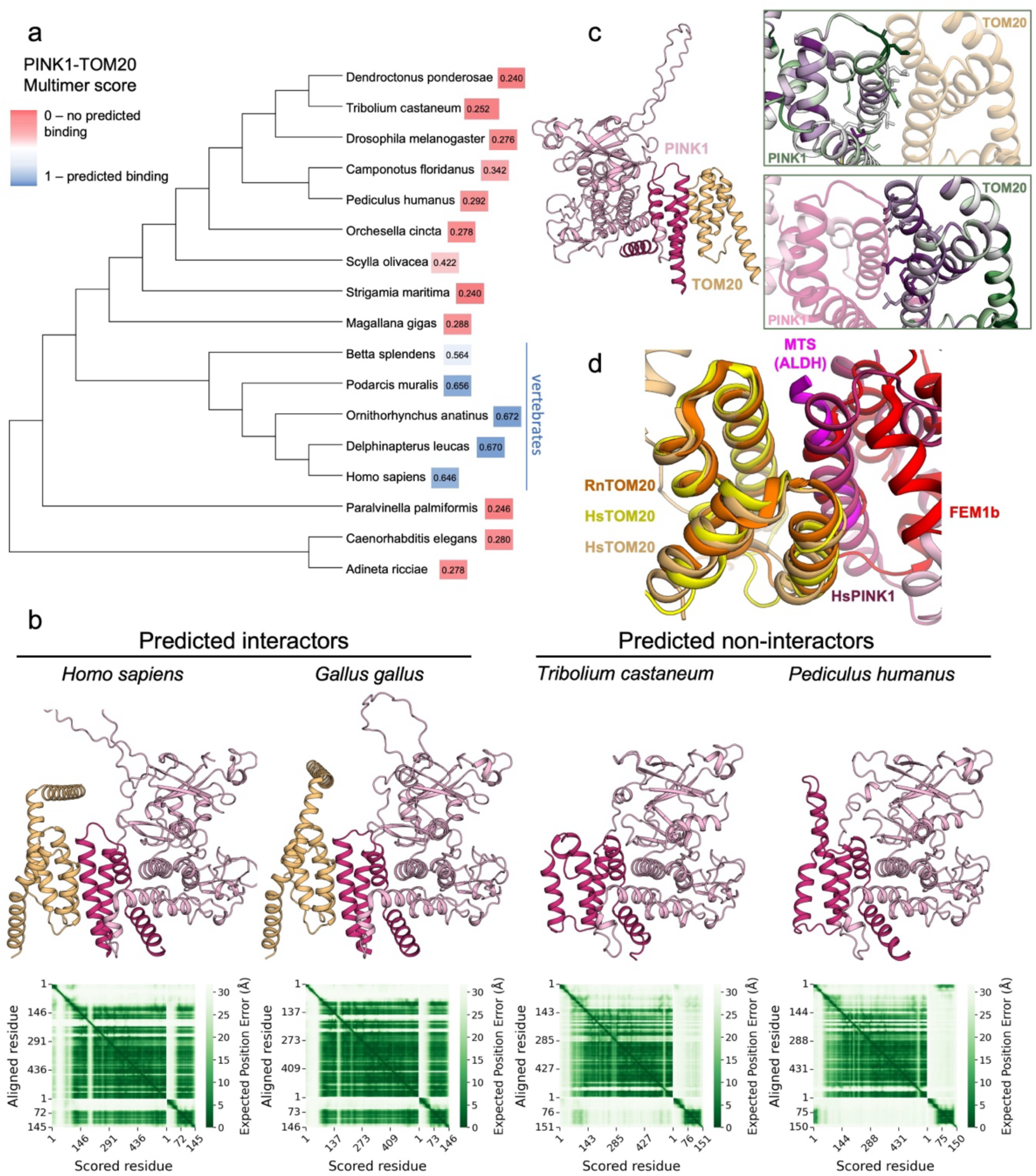
Vertebrate-specific evolution of the PINK1–TOM20 interaction. **(a)** Phylogenetic tree of representative metazoan PINK1 orthologues with corresponding AlphaFold3 multimer scores for the predicted PINK1–TOM20 complex. Scores range from 0 to 1, with higher values indicating greater confidence in the predicted complex. Vertebrate species are indicated with the blue bar. **(b)** Representative AlphaFold3 models and predicted aligned error (PAE) plots for PINK1–TOM20 complexes from two vertebrate predicted interactors (*Homo sapiens* and *Gallus gallus*) and two invertebrate predicted non-interactors (*Pediculus humanus* and *Tribolium castaneum*). PINK1 mitochondrial targeting sequences were omitted from the structural representations for clarity. **(c)** Evolutionary conservation of the PINK1–TOM20 interface. Residue conservation calculated from multiple sequence alignments of PINK1 and TOM20 orthologues was mapped onto the HsPINK1–HsTOM20 complex using ConSurf. The N-terminal region of PINK1 was omitted for clarity. **(d)** Structural comparison of the TOM20 interaction surface used by PINK1 and other TOM20-binding proteins. Structures of TOM20 bound to PINK1, a mitochondrial targeting sequence (MTS), and FEM1B are shown (PDB IDs 9EIH, 2V1T, and 9JCE, respectively).

To determine whether this divergence arose from changes in PINK1 or the mitochondrial import machinery, we examined the conservation of the TOM complex components across the same species panel. In contrast to PINK1, TOM20 was highly conserved throughout vertebrates and invertebrates (Table S1, Suppl. Fig. S2). Notably, residues previously predicted to mediate the PINK1–TOM20 interaction were largely conserved within TOM20, whereas the corresponding interface on PINK1 was highly variable (Fig. 2c). This is consistent with TOM20 having to bind a large number of protein motifs, including hundreds of mitochondrial targeting sequences which bind to the same interface (Fig. 2d). This would subject TOM20 to a stronger evolutionary pressure for conservation to retain binding to all MTSs, whereas TOM20 recognition by PINK1 would represent a more recent evolutionary adaptation. Furthermore, a recent cryoEM structure of the CRL2–FEM1B–TOM20 complex identified residues 17–25 within the ANK1 repeat of FEM1B as the TOM20-binding interface (Raiff et al., 2026). Like PINK1, the TOM20-binding interface of FEM1B is the same that engages PINK1 (Fig. 2d). Together, these observations suggest that vertebrate-specific TOM20 recognition by PINK1 arose primarily through sequence changes within PINK1 rather than TOM20 itself.

### Human PINK1, but not *Tribolium castaneum* PINK1, directly binds TOM20 through the NTE-CTE region

Our AlphaFold3 analysis predicted that vertebrate-specific sequence changes within the PINK1 NTE-CTE region mediate TOM20 recognition. To test this prediction, we examined PINK1-TOM20 interactions using 2D H^1^-N^15^ HSQC NMR spectroscopy. We first produced recombinant ^15^N,^13^C-labeled HsTOM20 (soluble domain, a.a. 51-145), and assigned its backbone amide ^1^H-^15^N resonances using standard 3D triple resonance sequential assignment methods. This produced high-quality HSQC spectra with 98% complete assignment (Fig. 3a). Next, as recombinant HsPINK1 remains difficult to produce, we engineered a TcPINK1 variant in which the ten NTE-CTE residues at the TOM20 interface were substituted with the corresponding human residues (humanized variant called TcPINK1^HsNC^). A sequence alignment shows that these residues are not conserved between HsPINK1 and TcPINK1 (Fig. 3b) and AlphaFold3 predicts that the humanized TcPINK1 should bind to HsTOM20 (Suppl. Fig. S3a). Addition of unlabeled TcPINK1^HsNC^ to ^15^N-labeled TOM20 produced clear chemical shift perturbations or line broadening throughout the HSQC spectrum, demonstrating direct interaction between the engineered PINK1 construct and HsTOM20 (Fig. 3a, b, Suppl. Fig. S3b). The perturbations occurred primarily on helices 1, 2, and 3 of TOM20, and map to the interface that binds the PINK1 NTE-CTE (Fig. 3c, d). To determine whether these perturbations resulted specifically from the engineered interface, wild-type TcPINK1 was incubated with ^15^N-HsTOM20 and showed no significant perturbations (Fig. 3e). Furthermore, incubation of ^15^N-TcTOM20 with TcPINK1 also showed no interaction, confirming the prediction that PINK1 does not bind TOM20 in insects (Fig. 3f). Together, these findings provide direct biochemical validation of the AlphaFold3 predictions and demonstrate that introduction of only ten vertebrate-specific interface residues is sufficient to confer TOM20 recognition.

**Figure 3.**
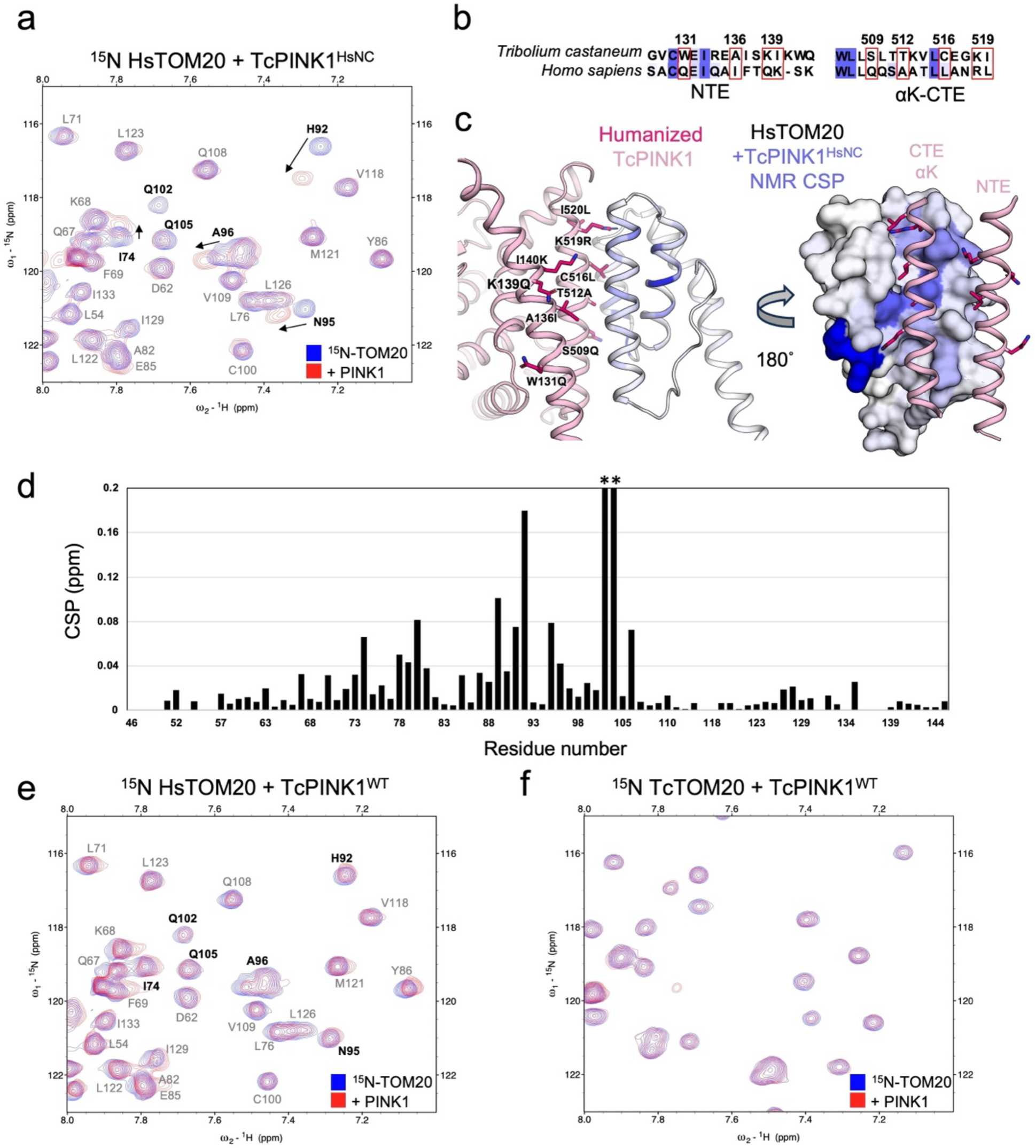
The PINK1 NTE–CTE interface determines TOM20 recognition. **(a)** Overlay of ^1^H–^15^N HSQC spectra of ^15^N-labelled HsTOM20 (51–145) in the absence and presence of TcPINK1 carrying ten human substitutions within the predicted TOM20-binding NTE–CTE interface (TcPINK1^HsNC^). **(b)** Sequence comparison of HsPINK1 and TcPINK1 highlighting the ten residues within the NTE–CTE TOM20-binding interface that were substituted with the corresponding human residues to generate TcPINK1^HsNC^. **(c)** Schematic and structural representation of TcPINK1^HsNC^. The ten residues at the predicted TOM20-binding interface substituted with their corresponding human residues are indicated. **(d)** Chemical shift perturbations (CSPs) of HsTOM20 residues following addition of TcPINK1^HsNC^. **(e)** ^1^H–^15^N HSQC analysis of ^15^N-labelled HsTOM20 in the absence and presence of wild-type TcPINK1, showing minimal perturbation compared with TcPINK1^HsNC^. **(f)** ^1^H–^15^N HSQC analysis of ^15^N-labelled TcTOM20 in the absence and presence of wild-type TcPINK1.

### Invertebrate PINK1 orthologues exhibit greater recombinant stability and activity

We next asked whether the evolutionary differences in TOM20 recognition were reflected in the stability and activity of recombinant PINK1 orthologues. We selected nine PINK1 orthologues, including 5 invertebrates and 4 vertebrates covering a large range of sequences (Suppl. Fig. S4), and expressed them as N-terminal GST fusion proteins in *E. coli*, followed by identical purification conditions. Analysis of purified protein revealed a striking difference between vertebrate and invertebrate PINK1 orthologues. All five invertebrate proteins yielded abundant and pure full-length recombinant protein following affinity purification, whereas vertebrate orthologues consistently produced substantially lower yields, with some evidence of degradation, despite identical cloning, expression, and purification conditions (Fig. 4a). To determine whether the purified proteins retained catalytic activity, GST-tagged PINK1 orthologues were incubated with tetra-ubiquitin substrate (Ub_4_), and ubiquitin phosphorylation was assessed by immunoblotting for phospho-ubiquitin. All five invertebrate orthologues robustly phosphorylated ubiquitin, whereas none of the vertebrate orthologues generated detectable phospho-ubiquitin signal under identical assay conditions (Fig. 4a). Thus, the ability to express recombinant PINK1 in bacteria closely paralleled its catalytic activity. The recombinant behavior of all nine orthologues also closely mirrored the AlphaFold3 predictions of their interaction with TOM20. Vertebrate PINK1 proteins, which were predicted to interact strongly with TOM20, consistently exhibited poor recombinant yield and lacked detectable kinase activity, whereas invertebrate orthologues with low predicted TOM20 interaction scores readily expressed as catalytically active kinases. Together, these findings reveal an evolutionary transition linking predicted TOM20 recognition with stability.

**Figure 4.**
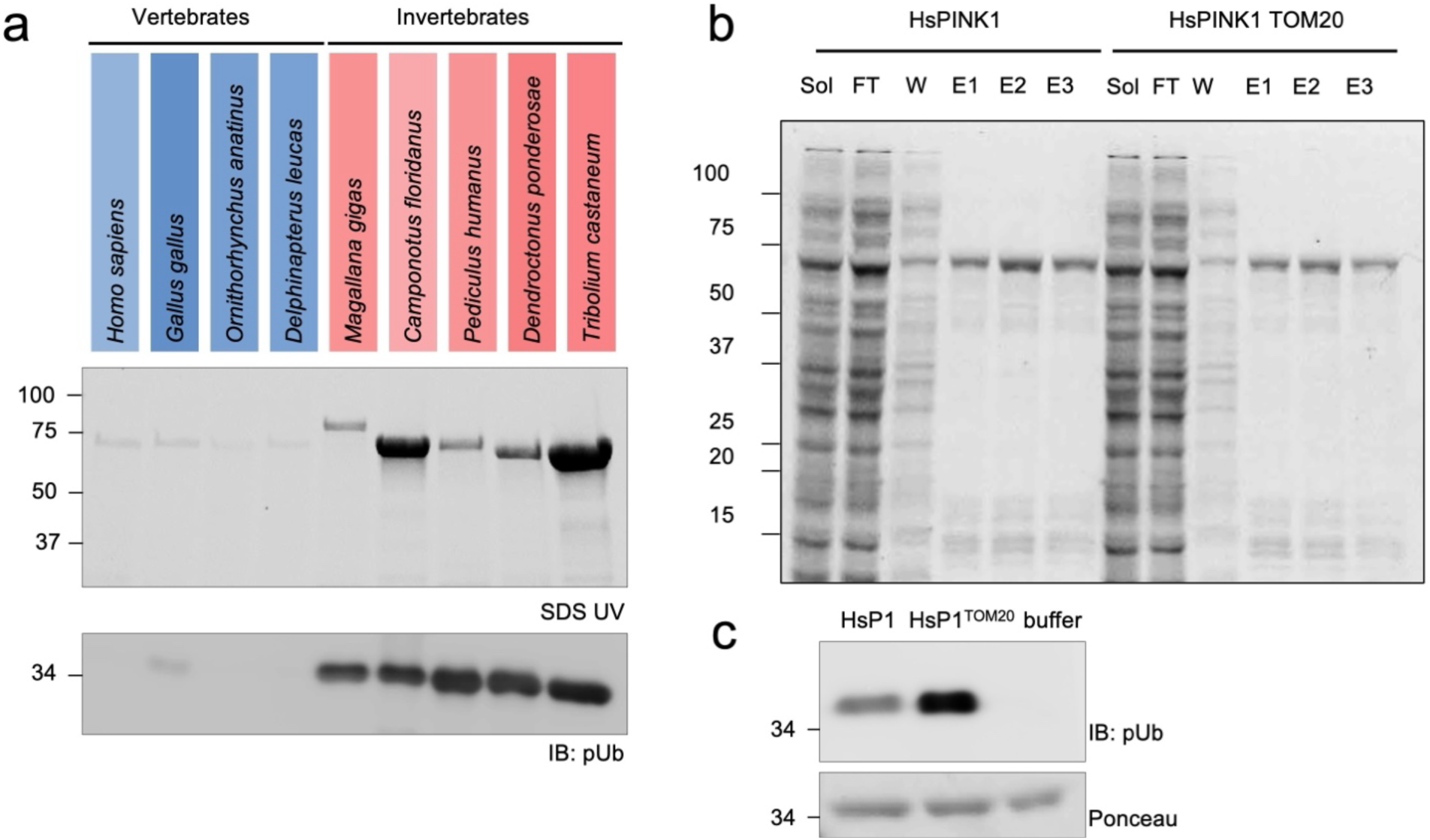
Recombinant activity of PINK1 orthologues correlates with evolutionary divergence in TOM20 recognition. **(a)** Recombinant expression and kinase activity of four vertebrate and five invertebrate PINK1 orthologues expressed as N-terminal GST fusion proteins in *E. coli*. Purified proteins were analysed by SDS–PAGE, and kinase activity was assessed using tetra-ubiquitin (Ub4) as substrate followed by immunoblotting for Ser65-phosphorylated ubiquitin (pUb). Vertebrate and invertebrate orthologues are indicated. The colour scale reflects the corresponding AlphaFold3-predicted PINK1– TOM20 interaction score for each orthologue. **(b)** GST affinity purification of HsPINK1 expressed alone or co-expressed with HsTOM20. Soluble lysate (Sol), flow-through (FT), wash (W), and sequential glutathione elution fractions (E1–E3) are shown. **(c)** Ubiquitin phosphorylation activity of recombinant HsPINK1 expressed alone or co-expressed with HsTOM20 (51–145) in *E. coli*. Equal amounts of purified HsPINK1 were incubated with Ub4, and phosphorylation was detected by pUb immunoblotting. Ponceau staining is shown as a loading control.

Having established that vertebrate-specific sequence changes are sufficient to confer TOM20 binding (Fig. 3), we next asked whether this interaction influences recombinant HsPINK1 activity during bacterial expression. HsPINK1^112-581^ and HsTOM20^51-145^ were co-expressed in *E. coli* using a pET-DUET dual-expression vector. As a control, HsPINK1 was expressed from the identical vector containing an empty second multiple cloning site. Recombinant proteins were purified under identical conditions, and equal purification volumes were analyzed by SDS-PAGE. Co-expression with HsTOM20 did not noticeably alter recombinant HsPINK1 yield, as comparable amounts of purified protein were recovered under both conditions (Fig. 4b). Identical purification volumes were analyzed, and these results indicate that TOM20 does not substantially increase recombinant protein production. However, co-expression with HsTOM20 increased ubiquitin phosphorylation compared with HsPINK1 expressed alone (Fig. 4c), indicating that TOM20 increases the proportion of catalytically active HsPINK1 generated during bacterial expression.

### Evolution-guided engineering restores recombinant human PINK1 activity

Our evolutionary and biochemical analyses suggested that divergence within the NTE-CTE region underlies the inability of HsPINK1 to express as an active recombinant kinase. We therefore engineered a series of chimeric proteins between HsPINK1 and TcPINK1 to determine whether these regions can revert the instability of HsPINK1 (Fig. 5a). The first chimeric protein (chimera 1’) was designed by replacing the human NTE– CTE with the corresponding TcPINK1 sequences. Because the NTE-CTE is divergent and potentially makes incompatible contacts with the kinase C-lobe, we added 10 mutations at this interface on the C-lobe (called chimera 1) (Fig. 5b and Fig. S4). The insert-1 is also much longer in HsPINK1 than in TcPINK1, and we thus shortened this insert, which appears to play no significant role in cells (Kumar et al., 2017) but could adversely affect expression and stability. The CTE interacts with TOM20 primarily via the NTE and αK helix of the CTE, and thus we created another chimera with only these regions swapped from TcPINK1 (named chimera 2), along with a number of interface mutations within the CTE (Fig. 5b). These two chimeras were expressed in *E. coli* and their yield, degradation, and activity were compared to HsPINK1 and TcPINK1. TcPINK1 consistently yielded more recombinant protein than all other constructs, whereas HsPINK1 and the chimeras were recovered at comparable levels following purification, with noticeable degradation (Suppl. Fig. S5b). Limited proteolysis also showed that TcPINK1 was considerably more resistant (Suppl. Fig. S5c). After normalizing protein amounts, a ubiquitin phosphorylation assay was conducted. As expected, HsPINK1 failed to phosphorylate ubiquitin, whereas TcPINK1 exhibited robust kinase activity (Fig. 5c). Strikingly, although TcPINK1 had more activity, chimera 1 had a significant amount of activity, whereas chimera 2 did not. Critically, reverting the 10 interface mutations between the CTE and C-lobe abrogated the activity of chimera 1, demonstrating that replacement of the NTE-CTE module alone is insufficient for folding of native PINK1 kinase and that the interactions between the NTE-CTE and the kinase C-lobe are crucial.

**Figure 5.**
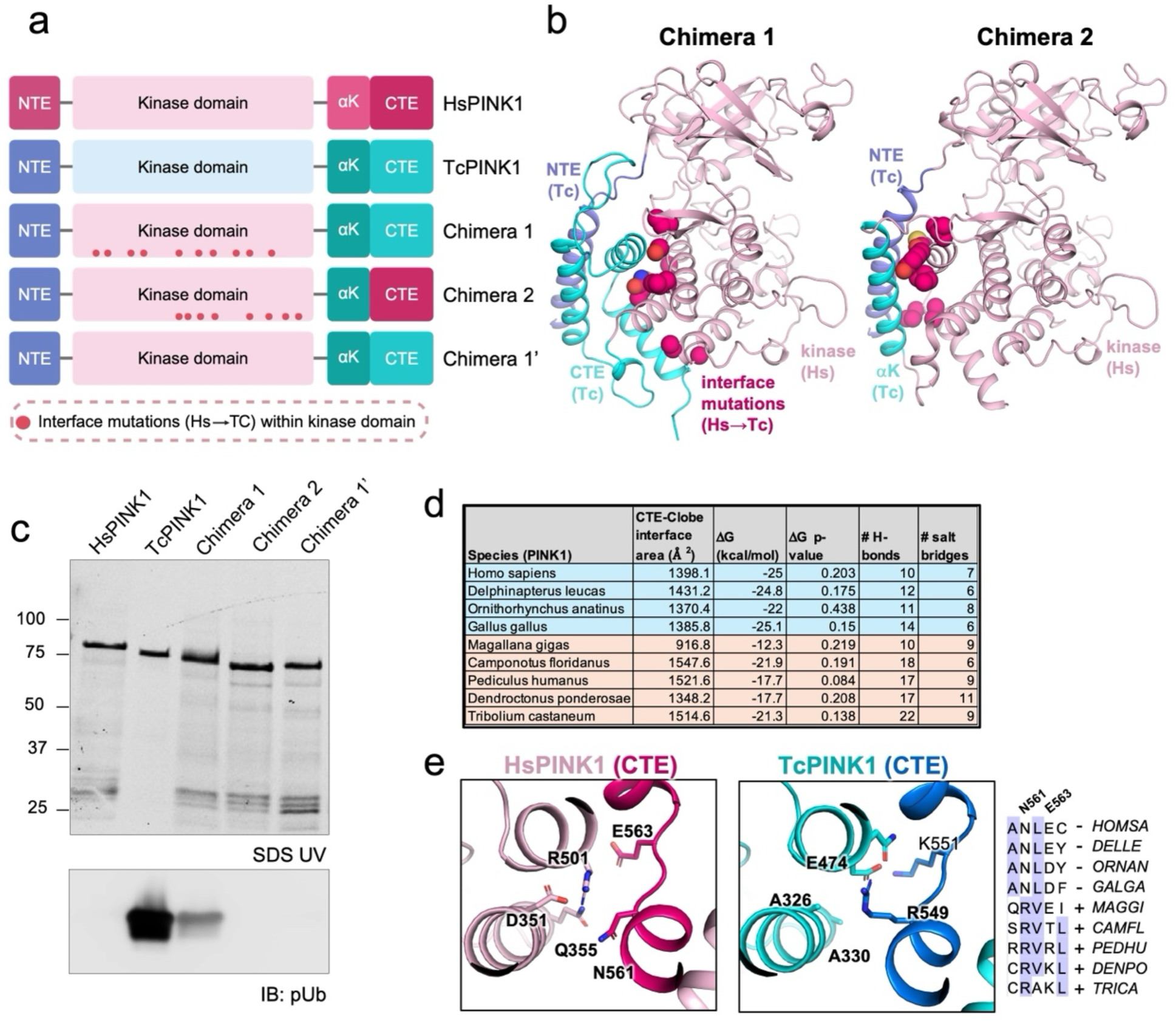
Evolution-guided engineering restores recombinant activity to the human PINK1 kinase domain. **(a)** Schematic representation of PINK1 chimeras generated by combining regions of HsPINK1 and TcPINK1. Chimera 1 contains the TcPINK1 NTE and CTE surrounding the HsPINK1 kinase domain together with ten TcPINK1-derived substitutions at the NTE–CTE–kinase interface. Chimera 2 contains the TcPINK1 NTE and αK region with the remaining kinase domain and CTE derived from HsPINK1. Chimera 1′ contains the same domain substitutions as Chimera 1 without the ten compensatory interface substitutions. **(b)** AlphaFold3 model of Chimera 1 highlighting the TcPINK1-derived NTE and CTE, HsPINK1 kinase domain, and the ten TcPINK1-derived interface substitutions. **(c)** Ubiquitin phosphorylation activity of recombinant HsPINK1, TcPINK1, Chimera 1, Chimera 2, and Chimera 1′. Protein amounts were normalized prior to incubation with Ub4, and phosphorylation was detected by pUb immunoblotting. Corresponding protein samples are shown by SDS–PAGE. **(d)** Structural analysis of the CTE–kinase C-lobe interface across the nine experimentally tested PINK1 orthologues. The number of predicted hydrogen bonds and salt bridges at the interface is shown for four vertebrate and five invertebrate PINK1 proteins. **(e)** Comparison of interactions at the CTE–C-lobe interface of HsPINK1 and TcPINK1. Selected residues contributing to hydrogen bonds and salt bridges are shown. In TcPINK1, Arg549 forms a salt bridge with Glu474 at the base of the CTE–C-lobe interface.

To understand how the CTE of PINK1 stabilizes the kinase domain, we performed an *in silico* structural analysis of its interface across the nine orthologues that we tested experimentally (Fig. 5d and Suppl. Fig. S6). We first used PISA to calculate the buried surface area between the CTE and C-lobe, which showed no clear pattern. Likewise, estimated DG of the interface did not show significant differences between vertebrates and invertebrates. However, the number of hydrogen bonds and salt bridges was significantly higher in invertebrates (*p* = 0.045). For instance, we observed a network of H-bonds and salt bridges at the base of the C-lobe-CTE interface which is very different between HsPINK1 and TcPINK1 (Fig. 5e). In TcPINK1 CTE, this region contains Arg549, which forms a salt bridge with Glu474 in the C-lobe, and is conserved in invertebrates, but not vertebrates. In summary, the CTE confers its stability to the kinase domain in invertebrates PINK1 and in vertebrates.

### Engineered human/insect PINK1 chimera recapitulates the activity of PINK1 mutations in cells and identifies R152W as a pathogenic variant

Having established that Chimera 1 retains catalytic properties of HsPINK1, we next asked whether it could serve as a biochemical model to investigate PINK1 variants that may be pathogenic or provide mechanistic insights. Recombinant HsPINK1 remains difficult to obtain in sufficient quantities therefore the engineered chimera provides an opportunity to directly characterize the molecular consequences of variants in vitro. To this end, we first tested a naturally occurring variant of unknown significance (VUS) in the PINK1 kinase domain that is not conserved in invertebrates, namely R152W. Arg152 is located in the loop immediately following the NTE, and its sidechain forms a salt bridge with Glu252 in the Alphafold3 model of PINK1 (Fig. 6a). MDSgene classification interprets the R152W variant as “probably pathogenic” as it was found as a compound heterozygote in an individual affected by PD (Kasten et al., 2018), but is assigned as a VUS in ClinVar. To test its activity, human HeLa PINK1-KO cells were transfected with PINK1-3HA wild type (WT) or variants and then treated with CCCP to depolarize mitochondria, which is known to activate PINK1 (Narendra et al., 2010). As expected, WT PINK1 accumulates with CCCP and induces ubiquitin phosphorylation (Fig. 6c). The R152W variant significantly reduced ubiquitin phosphorylation, consistent with it being probably pathogenic (Fig. 6c). To determine if the effect was intrinsic to PINK1, we introduced the R152W mutation in chimera 1 and tested Ub_4_ phosphorylation in vitro and found that the R152W variant strongly reduced activity compared to WT (Fig. 6d). The effect was specific to the incorporation of a Trp residue, as the synthetic mutation R152A did not affect activity in cells, and reduced to a lesser extent in vitro. Consistent with this last result, the E252K and E252A variants did not affect Ub phosphorylation in cells (Fig. 6c), suggesting that the observed salt bridge is not critical for maintaining PINK1’s activity. Of note, E252K is also a naturally occurring variant assigned as a VUS on ClinVar but is likely benign as it does not affect activity in cells. Intriguingly, while the E252K mutation did not affect the activity of chimera 1 in vitro (thus mirroring activity in cells), the E252A mutation dramatically increased Ub_4_ phosphorylation (Fig. 6d). Overall, these observations suggest that the loss of function observed in the R152W variant arises from the structural disruption introduced by the bulky aromatic residue, and not the loss of a positive charge.

**Figure 6.**
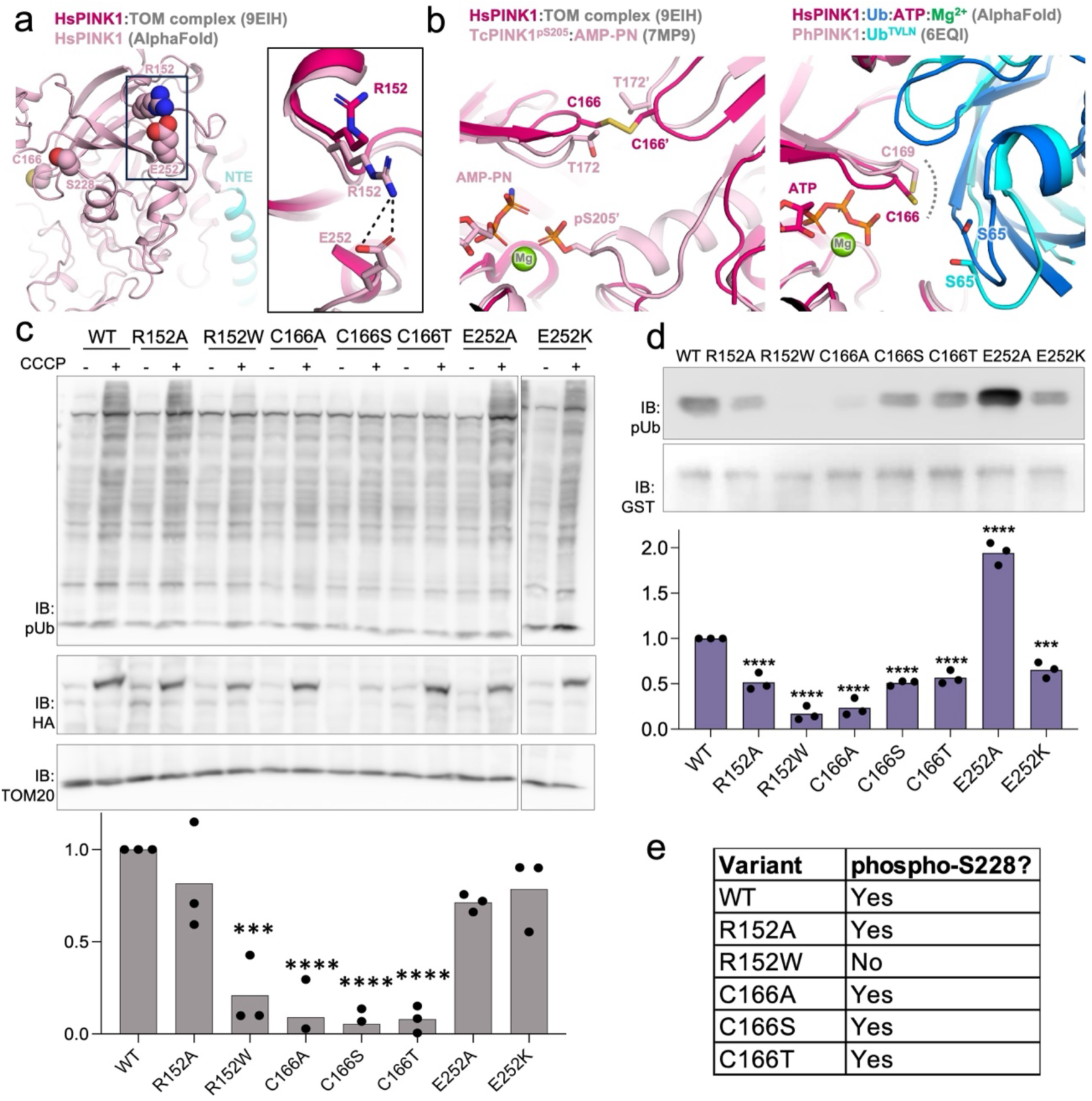
Impact of PINK1 mutations on autophosphorylation and ubiquitin phosphorylation. **(a)** Structural comparison of HsPINK1 in the TOM-bound structure (PDB 9EIH) with the AlphaFold model of HsPINK1. The inset shows the salt bridge between R152 and E252, indicated by the dashed line. **(b)** Structural comparison of HsPINK1 with TcPINK1 and PhPINK1. Left: Superposition of HsPINK1 from the TOM-bound structure and AMP-PNP-bound TcPINK1. Right: Superposition of an AlphaFold model of HsPINK1 bound to ubiquitin, ATP and Mg²⁺ with the PhPINK1–Ub^TVLN^ structure. Dotted lines indicate van der Waals interactions. **(c)** Cellular activity of WT HsPINK1 and the indicated R152, C166, and E252 variants in PINK1-knockout HeLa cells. Cells expressing HA-tagged PINK1 were treated with DMSO or 20 µM CCCP for 3 h. Lysates were assessed by immunoblotting for HA and Ser65-phosphorylated ubiquitin (pUb), respectively, with TOM20 shown as a loading control. CCCP-induced pUb was calculated by subtracting the corresponding DMSO signal and normalized to WT within each independent experiment (n=3). Statistical significance was assessed on the non-normalized CCCP − DMSO values using one-way ANOVA with Dunnett’s multiple-comparisons test against WT. *P < 0.05, **P < 0.01, ***P < 0.001, ****P < 0.0001. **(d)** Ubiquitin phosphorylation activity of recombinant WT Chimera 1 and the indicated R152, C166, and E252 variants. Following in vitro kinase assays with Ub4, pUb was detected by immunoblotting and normalized to GST-Chimera 1 (n=3). Statistical significance was assessed using one-way ANOVA with Dunnett’s multiple-comparisons test against WT Chimera 1. **(e)** Detection of Ser228 autophosphorylation in Chimera 1 variants by LC–MS. The table indicates whether a peptide containing phosphorylated Ser228 was detected for each variant.

To gain more insights into how variants alter PINK1’s activity, we further biochemically characterized these variants in Chimera 1. Because Chimera 1 precipitates when cleaved off from its tag, we kept them as fusion proteins with GST. Limited proteolysis shows a different pattern of digestion for R152W compared to WT, with a more pronounced band at 25 kDa in R152W, corresponding to free GST (Suppl. Fig. S7). This higher susceptibility to proteolysis suggests that the R152W variant may misfold. By contrast, the R152A and E252K variants show a pattern of digestion similar to WT, consistent with their level of activity being comparable to WT. Because these variants may also affect the ability of PINK1 to bind ubiquitin, without affecting the kinase activity proper, we next examined autophosphorylation. LC-MS analysis confirmed autophosphorylation of Ser228 in Chimera 1, demonstrating that the engineered kinase undergoes the activating phosphorylation event characteristic of PINK1 (Fig. 6e). A Ser228-phosphorylated peptide was not detected for R152W, whereas Ser228 phosphorylation was detected for R152A. These data overall suggest that the R152W variant affects the stability/folding of the kinase domain, which in turn reduces overall kinase activity, thus making the R152W variant very likely pathogenic.

Another aspect of PINK1 regulation that can be investigated with the chimera is the formation of disulfide bonds. Structural work on PINK1 from *Pediculus humanus corporis* (PhPINK1), as well as the cryoEM structure of human PINK1 bound to TOM, reveals a homotypic disulfide bond formed by a cysteine in the P-loop, which is Cys166 in human PINK1 (Fig. 6b) (Callegari et al., 2025; Gan et al., 2022). It was suggested that formation of this disulfide bond plays a role in the down regulation of PINK1 in the presence of reactive oxygen species (Gan et al., 2022). However, this residue is substituted to a threonine in TcPINK1, as well as to an alanine in another insect PINK1 orthologue (Fig. 6b and Suppl. Fig. S4). To investigate the functional importance of this residue, Cys166 was substituted with alanine, serine, or threonine, both for expression in human cells or in Chimera 1. The three mutants exhibited almost complete loss of activity in cells (Fig. 6c). In vitro, the C166A mutation in the chimera 1 led to a strong loss of ubiquitin kinase activity, whereas substitution with serine or threonine partially preserved ubiquitin kinase function (Fig. 6d). Strikingly, all three Cys166 mutations were autophosphorylated at Ser228 (Fig. 6e), implying that these substitutions in the P-loop did not impair the intrinsic kinase activity of PINK1. Furthermore, AlphaFold3 modelling of HsPINK1 with ubiquitin or the Parkin ubiquitin-like domain, as well as the structure of PhPINK1 bound to a ubiquitin variant (Schubert et al., 2017), show that Cys166 (or equivalent in PhPINK1) mediates contact with ubiquitin/Ubl that would be disrupted by mutation (Fig. 6b). These observations strongly suggest that mutations of Cys166 impair ubiquitin recognition and that maintenance of a polar side chain at this position is important for catalytic activity.

## Discussion

A central observation of this study is the striking correlation between the evolutionary emergence of TOM20 recognition and the biochemical properties of PINK1 orthologues. Across a broad phylogenetic panel, vertebrate PINK1 proteins were predicted to bind TOM20 with high confidence (Fig. 2), whereas invertebrate orthologues generally lacked this interaction. This divergence was mirrored experimentally: invertebrate PINK1 proteins were readily purified as active kinases, whereas vertebrate orthologues consistently displayed poor recombinant behaviour and little or no detectable activity (Fig. 4a). Although correlation alone does not establish causality, the ability of TOM20 to enhance human PINK1 activity (Fig. 4c) and the direct binding observed by NMR (Fig. 3) together support a functional relationship between TOM20 recognition and kinase competence. The structure of PINK1 bound to the TOM complex also shows that the PINK1 kinase domain makes extensive contact with TOM20, but also with TOM5 and TOM40 (Callegari et al., 2025). These findings raise the possibility that, during vertebrate evolution, PINK1 became increasingly dependent on components of the mitochondrial import machinery for efficient folding, stabilization, or maintenance of its active conformation.

Our data further suggest that this evolutionary transition is encoded within the NTE-CTE region. Despite limited sequence conservation between vertebrate and invertebrate orthologues, the CTE makes extensive contacts with the kinase C-lobe in all available structures and is critical to fold the kinase domain. This is consistent with the previous observation that deletion of a.a. 487-570 in TcPINK1 decreases general kinase activity (Woodroof et al., 2011). Similarly, restoration of activity in the engineered chimera required replacement of the human NTE-CTE module together with compensatory substitutions at the interface with the kinase core (Fig. 5). This interface is notably different between vertebrate and invertebrate PINK1 orthologues, with the latter generally containing a larger network of hydrogen bonds and salt bridges at this interface, a feature frequently associated with enhanced protein stability (Panja et al., 2015). Collectively, these observations argue that the NTE-CTE module functions as an integral structural element of PINK1. Rather than serving merely as regulatory appendages, the NTE and CTE appear to act as an intramolecular scaffold that stabilizes the active kinase architecture.

Our findings also indicate that the human PINK1 kinase domain can adopt a catalytically competent conformation when provided with an appropriate structural environment, and is not intrinsically unable to fold, even in *E. coli*. Although less stable than insect PINK1, our “chimera 1” construct, which retains a predominantly human kinase domain sequence (except at the C-lobe interface), phosphorylated ubiquitin and underwent autophosphorylation at Ser228 (Fig. 5c, 6e). Furthermore, our data together with observations from other groups suggest that the inability to produce active recombinant human PINK1 in *E. coli* is not caused by the absence of a specific set of chaperones that are only found in eukaryotic cells. Indeed, affinity purification of human PINK1 overexpressed in mammalian cells yields a protein in a partially folded state bound to HSP90 and its accessory subunits CDC37 or FKBP51 (Tian et al., 2025). Thus, although HSP90 activity is required to activate PINK1 (Fiesel et al., 2017), HSP90 alone is not sufficient to fold human PINK1. However, overexpression in the presence of mitochondrial uncouplers yields folded human PINK1 bound to the TOM complex, emphasizing that folding requires interactions with TOM (Callegari et al., 2025).

The last observation raises an important question: given that human cells constitutively express TOM20 and the other TOM subunits, why is mitochondrial depolarization necessary to fold PINK1? Here, it is important to recall that PINK1 has two mitochondrial targeting sequences (MTS) that drive import across the outer mitochondrial membrane (OMM) and inner mitochondrial membrane (IMM). Deletion of the first MTS (a.a. 2-34) indeed renders PINK1 constitutively active, in the absence of depolarization (Bayne et al., 2025; Okatsu et al., 2015). In polarized mitochondria, the MTS-driven translocation of PINK1’s N-terminus into the matrix brings the NTE to the IMM, where it is irreversibly cleaved by PARL (Jin et al., 2010). If the NTE is in the IMM, it cannot interact with TOM20 in the cytosol, an interaction that is critical for PINK1 activation (Eldeeb et al., 2024). Furthermore, comparison of the structures of PINK1 bound to HSP90 (partially folded) or TOM20 (folded) shows that the NTE in its folded position would clash with HSP90 and would be unable to bind TOM20 (Fig. 7a). Overall, these observations suggest a specific sequence to the folding process: 1) chaperoning of the kinase-CTE by HSP90, 2) retention of the NTE on the cytosolic side, 3) release of the C-lobe/CTE from HSP90, 4) binding of the NTE-CTE to TOM20 (Fig. 7b). The folded PINK1 kinase can then autophosphorylate in *trans* via another PINK1 bound to TOM and connected via a VDAC2 spacer (Callegari et al., 2025).

**Figure 7.**
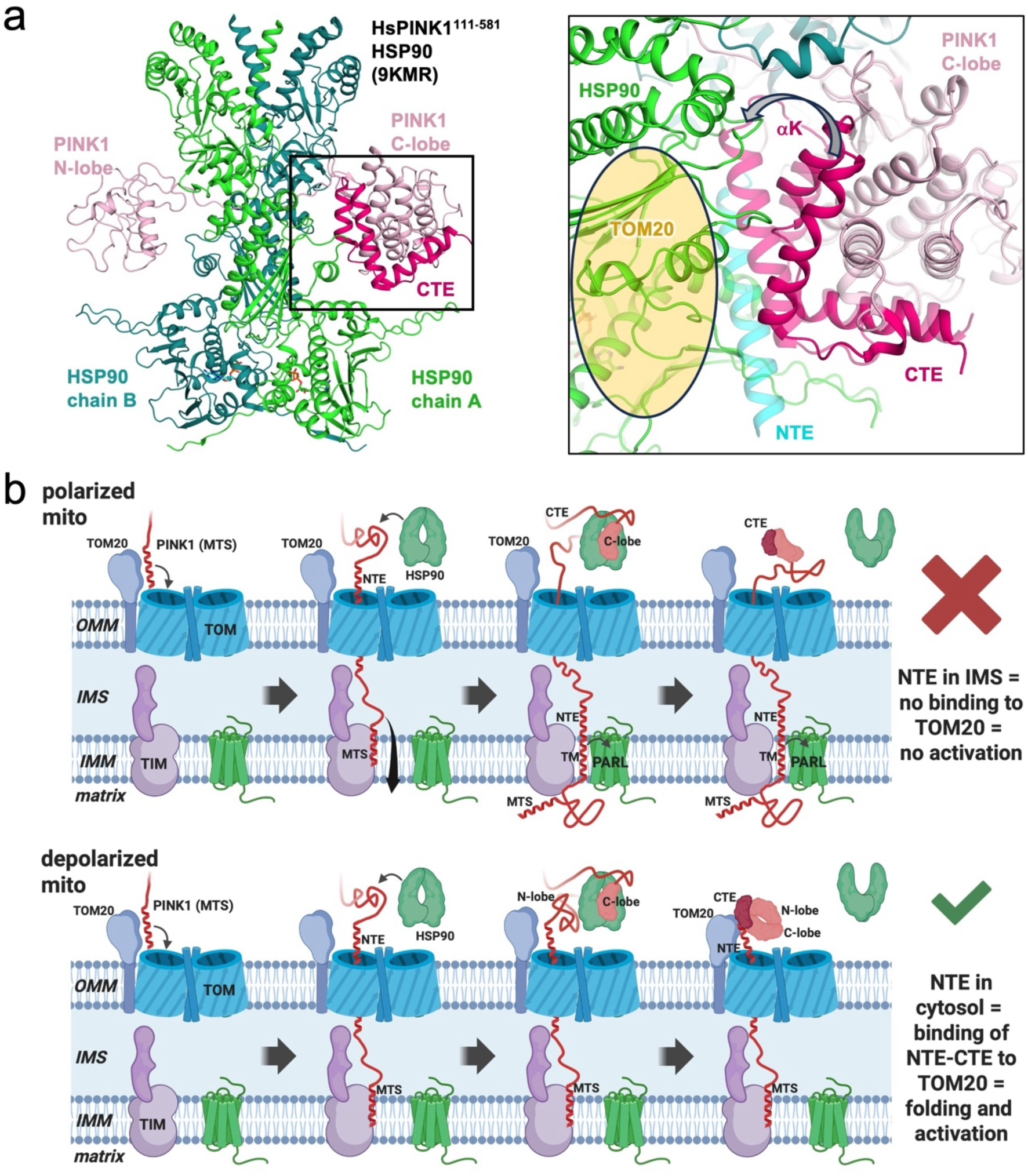
Model of PINK1 folding and activation. **(a)** Structure of human PINK1 (HsPINK1^111–581^) in complex with an HSP90 chaperone dimer (PDB 9KMR). The N-terminal extension (NTE), C-terminal extension (CTE), and αK helix are highlighted. The region proposed to interact with TOM20 is indicated and is inaccessible in the HSP90-bound conformation. **(b)** Proposed model for the role of mitochondrial import and TOM20 in PINK1 folding and activation. In polarized mitochondria (top), PINK1 is imported through the TOM complex and translocated across the inner mitochondrial membrane by the TIM complex. This positions the NTE within the intermembrane space, preventing productive engagement of the NTE/CTE region with TOM20 and thereby limiting formation of an active kinase conformation. Following mitochondrial depolarization (bottom), PINK1 import across the inner mitochondrial membrane is arrested, retaining the NTE on the cytosolic face of the outer mitochondrial membrane. HSP90 initially stabilizes cytosolic PINK1, followed by engagement of TOM20 with the exposed NTE/CTE region. We propose that this interaction promotes folding and/or stabilization of the kinase domain into a catalytically competent conformation, enabling PINK1 activation.

Our engineered human-insect PINK1 chimera also provides insight into the molecular basis of disease-associated variants and the general mechanism of activation. The variant R152W clearly impaired autophosphorylation, ubiquitin phosphorylation, and resistance to proteolysis, consistent with a defect in structural integrity rather than a selective disruption of substrate recognition or impairment of interactions mediated in cells. This variant had been investigated in a mitophagy assay by another group previously and found not to be significantly impaired (Ma et al., 2021). However, our results clearly show impairment, consistent with this variant being likely pathogenic. Arg152 is exposed to the solvent and mutation to a Trp does not induce clashes in either the AlphaFold or cryo-EM structure of human PINK1 (Fig. 6a). The R152A mutation being nearly as active as WT excludes the possibility that a salt bridge mediated by Arg152 is important. Thus, the specific R152W variant likely leads to misfolding. In contrast, mutations targeting Cys166 preserved autophosphorylation while strongly reducing ubiquitin phosphorylation, suggesting a direct role of Cys166 in ubiquitin substrate engagement, a conclusion supported by the structural models of PINK1 bound to ubiquitin. These observations highlight the utility of the chimera as a platform for distinguishing defects in folding, activation, and substrate recognition, analyses that have been difficult to perform using currently available recombinant systems.

In summary, our findings support a model in which vertebrate PINK1 evolved increased dependence on TOM20-associated stabilization mechanisms, concomitant with divergence of the NTE-CTE module that supports kinase-domain integrity. By exploiting these evolutionary differences, we generated an active chimeric kinase containing a predominantly human catalytic domain and revealed intramolecular interactions that are critical for PINK1 function. Beyond providing insight into the molecular basis of PINK1 activation, this work establishes a framework for investigating disease-associated variants and offers a broadly useful platform for future structural and therapeutic studies of this important Parkinson’s disease kinase.

## Methods

### Identification of PINK1 homologues and phylogenetic analysis

PINK1 homologues were identified across metazoan species using HMMER v3.4. A PINK1 profile hidden Markov model (HMM) was generated using hmmbuild and searched against a metazoan protein sequence dataset obtained from UniProtKB (taxonomy ID: 33208) using hmmsearch. Hits were filtered using a full-sequence E-value threshold of 1 × 10⁻²⁰, and sequences shorter than 300 amino acids were excluded to remove incomplete or truncated proteins. Candidate homologues were clustered using CD-HIT at 80% sequence identity to reduce redundancy, and annotated fragments were removed. Representative sequences were subsequently selected to preserve broad taxonomic coverage while minimizing redundancy, resulting in a final dataset of 48 PINK1 homologues. Sequences were aligned using Clustal Omega with default parameters. Phylogenetic relationships were inferred by maximum-likelihood analysis in MEGA12 using the LG amino-acid substitution model with gamma-distributed rate variation. Branch support was assessed using 1,000 adaptive bootstrap replicates.

### Identification of TOM20 homologues

TOM20 homologues corresponding to species included in the PINK1 dataset were identified using an HMM-based search with the human TOM20 sequence (UniProt Q15388) as a query. Candidate sequences were filtered to retain one representative per species based on annotation and HMMER score, prioritizing canonical TOM20 proteins and excluding TOM20-like proteins and fragments.

### Structural and evolutionary analyses

Evolutionary conservation of PINK1 and TOM20 residues was assessed using the ConSurf server (Yariv et al., 2023). Conservation scores were calculated using the multiple sequence alignment of the 48 PINK1 and TOM20 orthologues generated as described above and mapped onto PINK1 structures for visualization. Protein interfaces were analyzed using the Protein Interfaces, Surfaces and Assemblies (PISA) server. PINK1 structures were submitted to PISA to characterize interactions between the C-terminal extension (CTE) and kinase domain. Interface properties, including buried surface area, predicted solvation free-energy gain upon interface formation (ΔG), hydrogen bonds, and salt bridges, were obtained from the PISA interface analysis and compared between PINK1 structures.

### AlphaFold3 multimer modeling and interaction scoring

Predicted interactions between PINK1 and TOM20 were assessed using AlphaFold3 through the AlphaFold Server (Abramson et al., 2024). For each of the 48 species included in the comparative analysis, full-length PINK1 and the corresponding TOM20 sequence were provided as input, and PINK1–TOM20 complex models were generated using default server settings. Model confidence was assessed using the predicted template modelling score (pTM) and interface predicted template modelling score (ipTM). A combined ranking score was calculated as 0.8 × ipTM + 0.2 × pTM (Evans et al, 2022). Ranking scores were compared across species to assess differences in predicted PINK1–TOM20 interaction confidence between vertebrate and invertebrate orthologues.

### PINK1 and TOM20 constructs design

All recombinant PINK1 constructs were synthesized by GenScript, codon-optimized for expression in Escherichia coli, and cloned into pGEX-6P-1 for expression as N-terminal GST fusion proteins. *Homo sapiens* PINK1 (HsPINK1) comprised residues 112–581, whereas *Tribolium castaneum* PINK1 (TcPINK1) comprised residues 121–570, as previously described (Rasool et al., 2018). Other PINK1 orthologs: *Gallus gallus* (chicken) a.a. 81-546; *Ornithorhynchus anatinus* (platypus) a.a. 93-558; *Delphinapterus leucas* (beluga whale) a.a. 118-584; *Magallana gigas* (Pacific oyster) a.a. 119-600; *Camponotus floridanus* (carpenter ant) a.a. 124-612; *Pediculus humanus corporis* (body louse) a.a.118-575; *Dendroctonus ponderosae* (mountain pine beetle) a.a. 123-570. Chimeric PINK1 constructs were designed by combining regions of HsPINK1 and TcPINK1 based on sequence alignment and structural modelling. Chimera 1 contained the TcPINK1 N-terminal region spanning residues 121–146, followed by HsPINK1 residues 142–183 and the human kinase domain spanning residues 213–513, with insert 1 deleted. The C-terminal extension (CTE) was replaced with TcPINK1 residues 487–570. In addition, ten substitutions were introduced at the predicted interface between the TcPINK1 NTE–CTE module and the human kinase domain. Chimera 1′ contained the same domain composition as Chimera 1 but lacked the ten interface substitutions. Chimera 2 contained the TcPINK1 N-terminal region spanning residues 121–146 followed by HsPINK1 residues 142–523, with insert 1 deleted. Within the CTE, only the TcPINK1 αK-containing region spanning residues 500–515 was introduced, while the remaining C-terminal sequence was derived from HsPINK1. In addition, the TcPINK1-derived αK region and adjacent CTE interface contained the corresponding TcPINK1-specific substitutions introduced to preserve local interactions with the kinase C-lobe. The soluble domain of *Homo sapiens* TOM20 (a.a. 51-145) or *Tribolium castaneum* TOM20 (a.a. 56-151) were also synthesized with codon optimization and cloned in pGEX-6p1. For PINK1–TOM20 co-expression experiments, a pET-Duet dual-expression construct was generated containing GST-tagged HsPINK1(112–581) in multiple cloning site 1 (MCS1) and HsTOM20(51–145) in multiple cloning site 2 (MCS2). A corresponding control construct contained GST-HsPINK1(112–581) in MCS1 with MCS2 left empty.

### Mutagenesis

Point mutations were introduced into the indicated PINK1 constructs by PCR-based site-directed mutagenesis using mutagenic primers. PCR products were treated with DpnI to remove the parental plasmid and transformed into *E. coli*. Plasmid DNA was isolated from individual colonies through miniprep, and the presence of the desired mutation was confirmed by Sanger sequencing.

### Bacterial growth and induction of expression for PINK1

Plasmids were transformed into E. coli BL21(DE3) cells for recombinant protein expression. Three colonies of the transformation were used to inoculate a 10 mL starter culture of Lysogeny Broth (LB) with 0.1 mg/mL ampicillin, and the cultures were incubated at 37 °C, shaking at 180 rpm overnight. The starter cultures were added to either 300 mL or 1L of LB supplemented with 0.1 mg/ml of ampicillin and left to grow at 37 °C until the optical density at the 600 nm wavelength (OD600) reached between 0.8-1. Cultures were cooled down to 16 °C and 300 µM of IPTG was added to induce protein expression. The cultures were left overnight at 16 °C, shaking at 180 rpm. Cells were harvested by centrifugation at 3500 rpm for 30 min at 4 °C and the pellets were resuspended in 9 mL or 30 mL lysis buffer (0.025 mg/mL DNAse, 5 mM MgCl_2_, 1 mM PMSF, 0.1 mg/Ml lysozyme, 0.2% Tween 20, 10% glycerol, 0.1% Chaps, in 300 mM NaCl and 50 mM Tris– HCl pH 8) depending on the initial volume of growth. The resuspended pellets were sonicated at 4 °C for 20 s on–off intervals repeated 8 times.

### Expression of ^15^N and ^15^N,^13^C-labeled TOM20

Plasmids were transformed into E. coli BL21(DE3) cells for recombinant protein expression. One colony of the transformation was used to inoculate a 10 mL starter culture of Lysogeny Broth (LB) with 0.1 mg/mL ampicillin and the cultures were incubated at 37 °C, shaking at 180 rpm overnight. 5 mL of the starter culture were harvested and added to 1 L of minimal M9 media supplemented with 0.1 mg/ml ampicillin, ^15^NH_4_Cl, and either unlabelled D-glucose or ^13^C_6_-D-glucose (Cambridge Isotope Laboratories). Cells were grown at 37 °C until OD600 reached 0.6. Cultures were cooled down to 20 °C and 500 µM of IPTG was added to induce protein expression overnight. Cells were harvested by centrifugation at 3500 rpm for 30 min at 4 °C and the pellets were resuspended in 30 mL lysis buffer (0.025 mg/mL DNAse, 5 mM MgCl2, 1 mM PMSF, 0.1 mg/mL lysozyme, in 300 mM NaCl and 50 mM Tris–HCl pH 8) per liter of initial volume of growth. The resuspended pellets were sonicated at 4 °C for 20 seconds on, 40 seconds off intervals for a total of 2:40 minutes on.

### Bacterial cell lysis and protein purification

Following cell lysis and sonication, the bacteria were spun down at 15,000 rpm for 30 minutes at 4 °C into a soluble fraction and insoluble pellet. The clarified cell lysate (supernatant) was incubated with 2 mL Sepharose 4B resin (GE Healthcare Life Sciences) resuspended in 2mL kinase buffer (50 mM Tris, 300 mM NaCl, 3 mM DTT, pH 8.0) for 1 h at 4 °C on a rotating platform. Sample was then transferred onto a gravity filtration column and the flow through fraction was saved. The bound resin was washed repeatedly with kinase buffer (wash fractions), and the protein of interest was eluted with 20 mM glutathione (Bio Basics Canada) and concentrated by centrifugation using 15 kDa Centrifugal Filter Units (Millipore) according to the manufacturer’s instructions. The protein concentration was measured on the Denovix DS-11 spectrophotometer at 280 nm wavelength. For NMR samples (TcPINK1, TcPINK1^HsNC^ and ^15^N,^13^C-labeled TOM20), proteins were cleaved overnight at 4 °C with 3C protease (1:50 ratio) and then further purified by size-exclusion chromatography on a Cytiva Superdex 75 (TOM20) or Superdex 200 (PINK1) column equilibrated in HBS buffer (20 mM HEPES, 150 mM NaCl, 3 mM DTT, pH 7.4).

### NMR

All datasets were acquired at 298K. Backbone assignments were performed on ^15^N,^13^C-labeled human TOM20^51-145^ at 1 mM in 10% D_2_O and HBS buffer. Gradient-enhanced CBCACONH, HNCACB, and HNCO experiments (Klukowski et al., 2023) were acquired pn 800 MHz Bruker Avance NMR spectrometer equipped with a triple-resonance (^1^H,^13^C,^15^N) cryoprobe. NMR spectra were processed using TopSpin 3.6.2. Resonances were automatically using the online platform NMRtist (Klukowski et al., 2023). 100% of backbone ^15^N-^1^H resonances were assigned and considered “strong” for accuracy. Titration experiments were performed using ^15^N-labeled HsTOM20 or TcTOM20 at 0.2 mM in 10% D_2_O and HBS buffer. Sensitivity-enhanced echo/anti-echo ^15^N-^1^H HSQC (Schleucher et al., 1994) were acquired on a 600 MHz Bruker Avance NMR spectrometer equipped with a triple-resonance (^1^H,^13^C,^15^N) cryoprobe. Spectra were analyzed using Poky (Chiu & Lee, 2026). Chemical shift perturbations were calculated using the formula 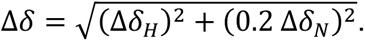

### Kinase assays

Kinase activity of purified recombinant PINK1 proteins was assessed using tetra-ubiquitin (Ub4) as substrate, based on a previously described assay (Rasool *et al*., 2024). Purified GST-tagged PINK1 proteins were normalized to a final concentration of 1 µM and incubated with approximately 3 µM tetra-ubiquitin (Ub4) in a 200 µL reaction containing 50 mM Tris-HCl, 300 mM NaCl, 3 mM DTT, pH 8.0, 100 µM ATP and 5 mM MgCl₂. Reactions were incubated at 30 °C for 10 min and terminated by addition of Laemmli sample buffer. Samples were resolved by SDS–PAGE and transferred to PVDF membranes. Ser65-phosphorylated ubiquitin was detected by immunoblotting using an anti-pSer65-Ub antibody. pUb signal was quantified by densitometry using ImageLab and normalized as indicated for each experiment.

### Limited proteolysis

Purified recombinant PINK1 proteins (10 µg) were subjected to limited proteolysis using elastase to assess differences in protease susceptibility. Proteins were incubated with elastase at protease dilutions of 1:800, 1:400, and 1:200 for 10 min at 37 °C. Reactions were terminated by addition of Laemmli sample buffer, and samples were resolved by SDS–PAGE. Proteolytic profiles were visualized by Coomassie staining.

### Trypsin digest and mass spectrometry

Protein samples were denatured in 6 M urea, 1 mM EDTA, and 50 mM triethylammonium bicarbonate (TEAB), pH 8.5, and digested with trypsin at an enzyme to protein ratio of 1:100 overnight at 37 °C. Following digestion, cysteine residues were reduced with 5 mM tris(2-carboxyethyl)phosphine (TCEP) for 10 min at 37 °C and subsequently alkylated with 50 mM iodoacetamide for 30 min at room temperature in the dark. Digested peptides were purified using C18 spin columns (Thermo Fisher Scientific) according to the manufacturer’s instructions.

Peptides were eluted in 2% formic acid and 0.5% acetonitrile, and 1 µg of peptide was loaded onto an Acclaim PepMap 100 C18 column. Peptides were separated using a 30-min gradient from 5–40% acetonitrile in 0.1% formic acid at a flow rate of 300 nL/min. Eluted peptides were analyzed using a Bruker Q-TOF mass spectrometer equipped with an Apollo II ion-funnel electrospray ionization source. MS/MS data were analyzed using MaxQuant v2.7.5.0, with carbamidomethylation of cysteine specified as a fixed modification and oxidation of methionine and phosphorylation of serine, threonine and tyrosine residues specified as variable modifications.

### Cell culture

HeLa PINK1-knockout (PINK1-KO) cells were maintained in Dulbecco’s modified Eagle medium (DMEM) supplemented with 10% fetal bovine serum (FBS) and penicillin– streptomycin at 37 °C in a humidified atmosphere containing 5% CO₂. Cells were transfected with pcDNA3.1 with 5’ and 3’ UTR plasmids for attenuated expression encoding C-terminally 3×HA-tagged human PINK1 or the indicated PINK1 variants using polyethyleneimine (PEI), with 7 µg of plasmid DNA per transfection. Cells were incubated for 72 h following transfection and subsequently treated with either 20 µM CCCP to induce mitochondrial depolarization or DMSO as a vehicle control for 3 h prior to harvest.

### Cell lysis and immunoblotting

Following treatment, cells were washed with ice-cold phosphate-buffered saline (PBS) and lysed in RIPA buffer supplemented with 1% Triton X-100 and protease inhibitor. Lysates were incubated on ice for 30 min with vortexing every 10 min and subsequently clarified by centrifugation at 13,000 rpm for 30 min at 4 °C. Protein concentrations of the resulting supernatants were determined using a bicinchoninic acid (BCA) protein assay. Eight micrograms of total protein per sample were resolved by SDS–PAGE and transferred onto polyvinylidene fluoride (PVDF) membranes.

Membranes were blocked with 5% bovine serum albumin (BSA) in Tris-buffered saline containing 0.1% Tween-20 (TBST) for 1 h and incubated with primary antibodies against Ser65-phosphorylated ubiquitin (pUb) or HA, each at a 1:2,000 dilution. Membranes were washed with TBST and incubated with the appropriate HRP-conjugated secondary antibodies. Protein signals were detected using enhanced chemiluminescence and imaged using an ImageQuant LAS 500 imaging system (GE Healthcare).

### Quantification and statistical analysis of PINK1 kinase activity

For cell-based PINK1 activity assays, phospho-ubiquitin (pSer65-Ub) signal was quantified by densitometry using ImageLab. For each PINK1 construct, the CCCP-induced pUb response was calculated as CCCP pUb − DMSO pUb. Values were not normalized to PINK1-HA abundance, as changes in PINK1 accumulation or stability caused by the introduced mutations were considered part of their overall functional effect on PINK1 signalling. For graphical presentation, the CCCP-induced pUb response was normalized to the wild-type PINK1 response within each independent experiment. Statistical analyses were performed on the non-normalized CCCP pUb − DMSO pUb values from independent experiments using a one-way ANOVA with Dunnett’s multiple-comparisons test against wild-type PINK1.

For recombinant PINK1 kinase assays, pSer65-Ub and GST-PINK1 signals were quantified by densitometry using ImageLab. Ubiquitin phosphorylation activity was calculated as the pUb signal divided by the corresponding GST-PINK1 signal to account for differences in the amount of recombinant kinase present in each reaction. Values were normalized to the indicated control for graphical presentation. Statistical analyses were performed as indicated in the corresponding figure legends.

## Acknowledgments

We thank Jimmy Ibarra for assistance with computational analyses and programming. We thank Mark Hancock (SPR-MS Facility), Tara Sprules (Quebec/Eastern Canada High-Field NMR facility), and Kim Munro (CRBS Facility) for assistance with mass spectrometry, NMR spectroscopy, and technical support, respectively. Jean-François Trempe is a member of the Centre de Recherche en Biologie Structurale, a research centre funded by the Fonds de Recherche du Québec (health sector), grant #288558. This work was supported by a grant from the Canadian Institutes of Health Research, grant # PJT-186189.

## Author contributions

T.S. and J.-F.T. conceptualization; T.S., S.V., and J.-F.T. investigation; T.S. and J.-F.T. writing–original draft; T.S. and J.-F.T. writing–review & editing; T.S., S.V., and J.-F.T. visualization; J.-F.T. supervision; J.-F.T. funding acquisition.

