## Supplemental Figures for "Evolutionary analysis of PINK1 reveals key roles of the N- and C-terminal extensions and TOM20 in folding its kinase domain"

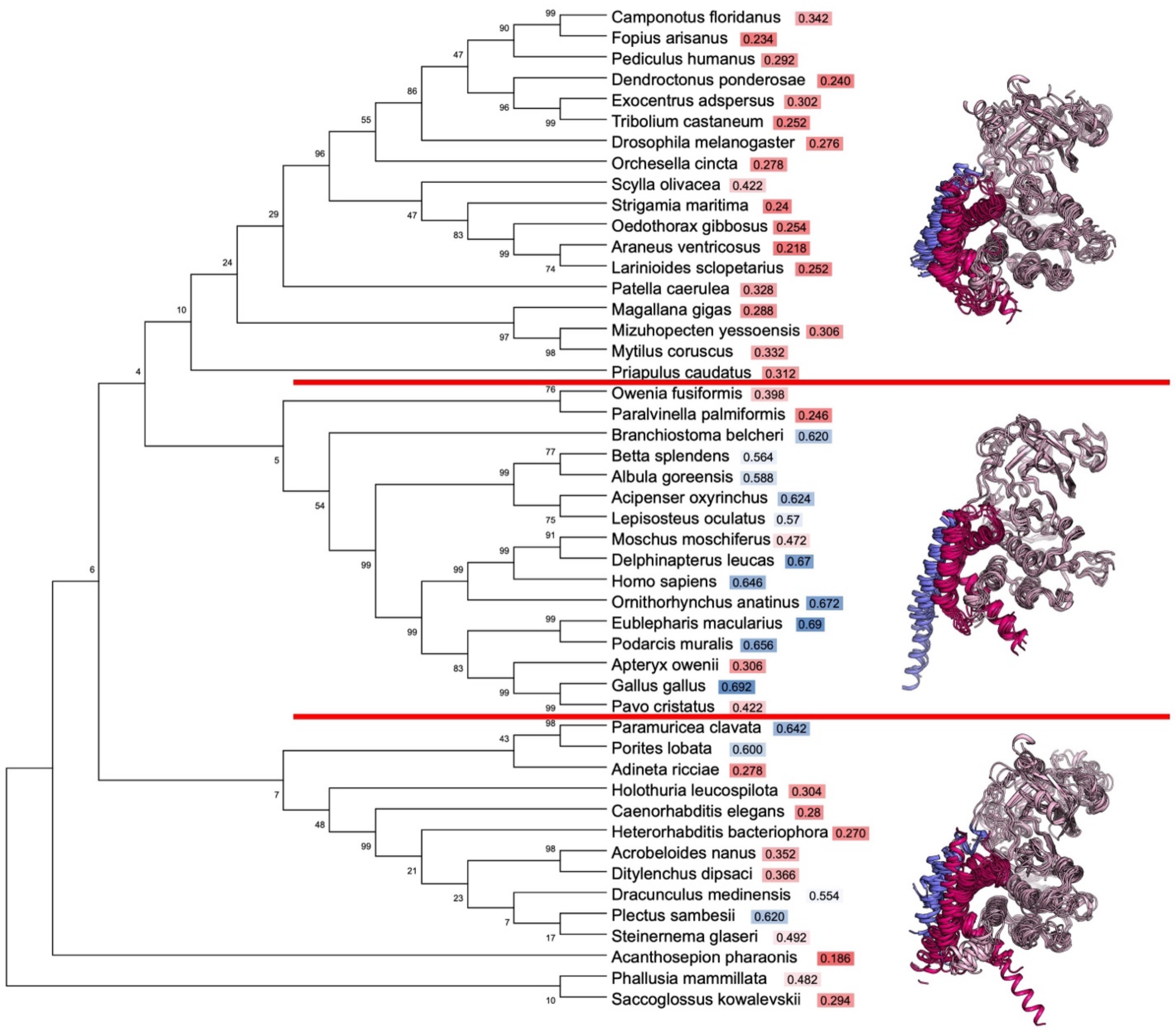


**Figure S1. Phylogenetic distribution of PINK1 orthologues used for evolutionary analysis**. Maximum-likelihood phylogenetic tree of 48 PINK1 orthologues used for evolutionary and AlphaFold3 analyses. The dataset spans representative vertebrate and invertebrate metazoan lineages.

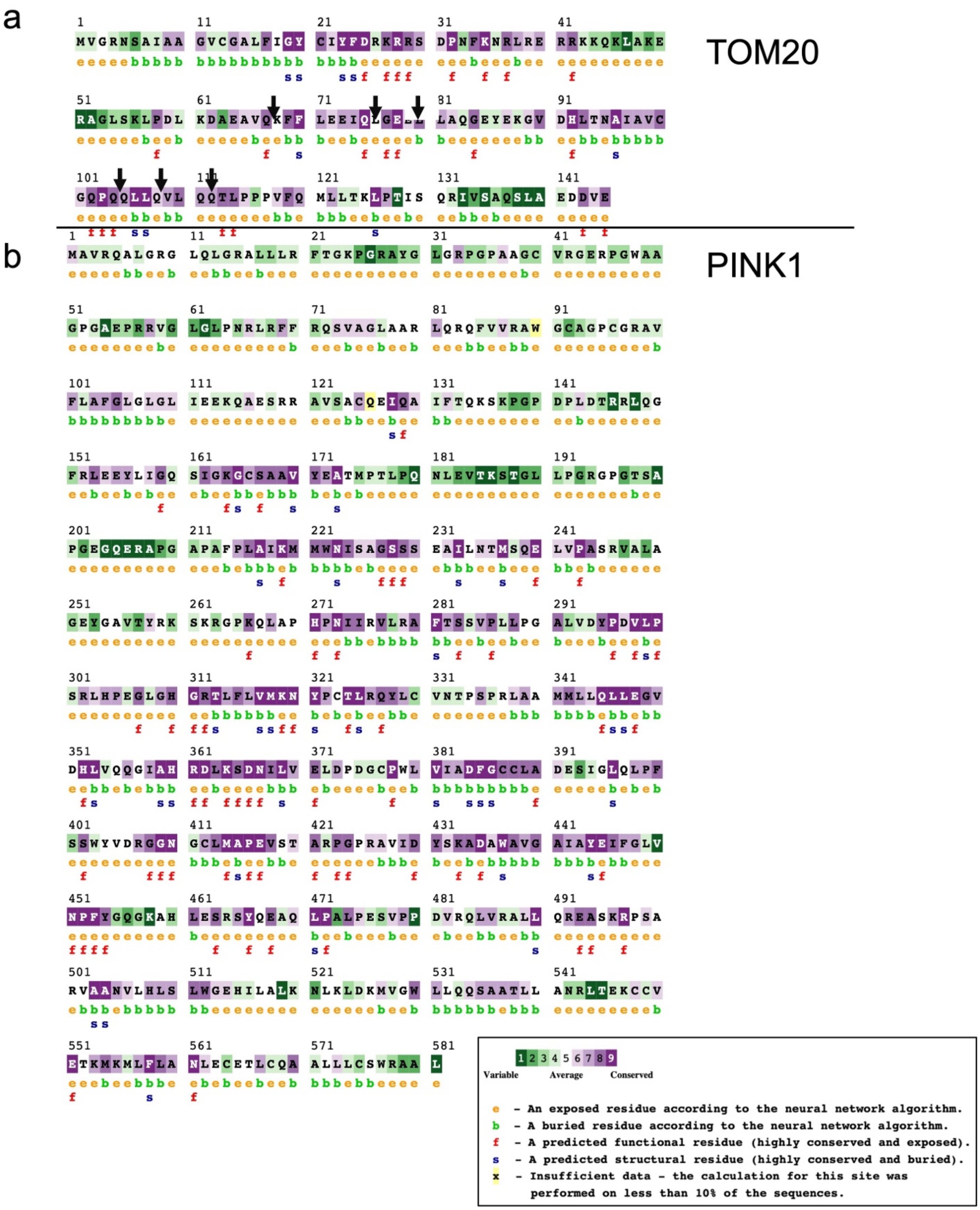


**Figure S2. Evolutionary conservation of TOM20 and PINK1**. **(a)** Evolutionary conservation of TOM20 through multiple sequence alignment across the analysed metazoan species using ConSurf. **(b)** Corresponding evolutionary conservation through multiple sequence alignment of PINK1 across the same species panel using ConSurf.

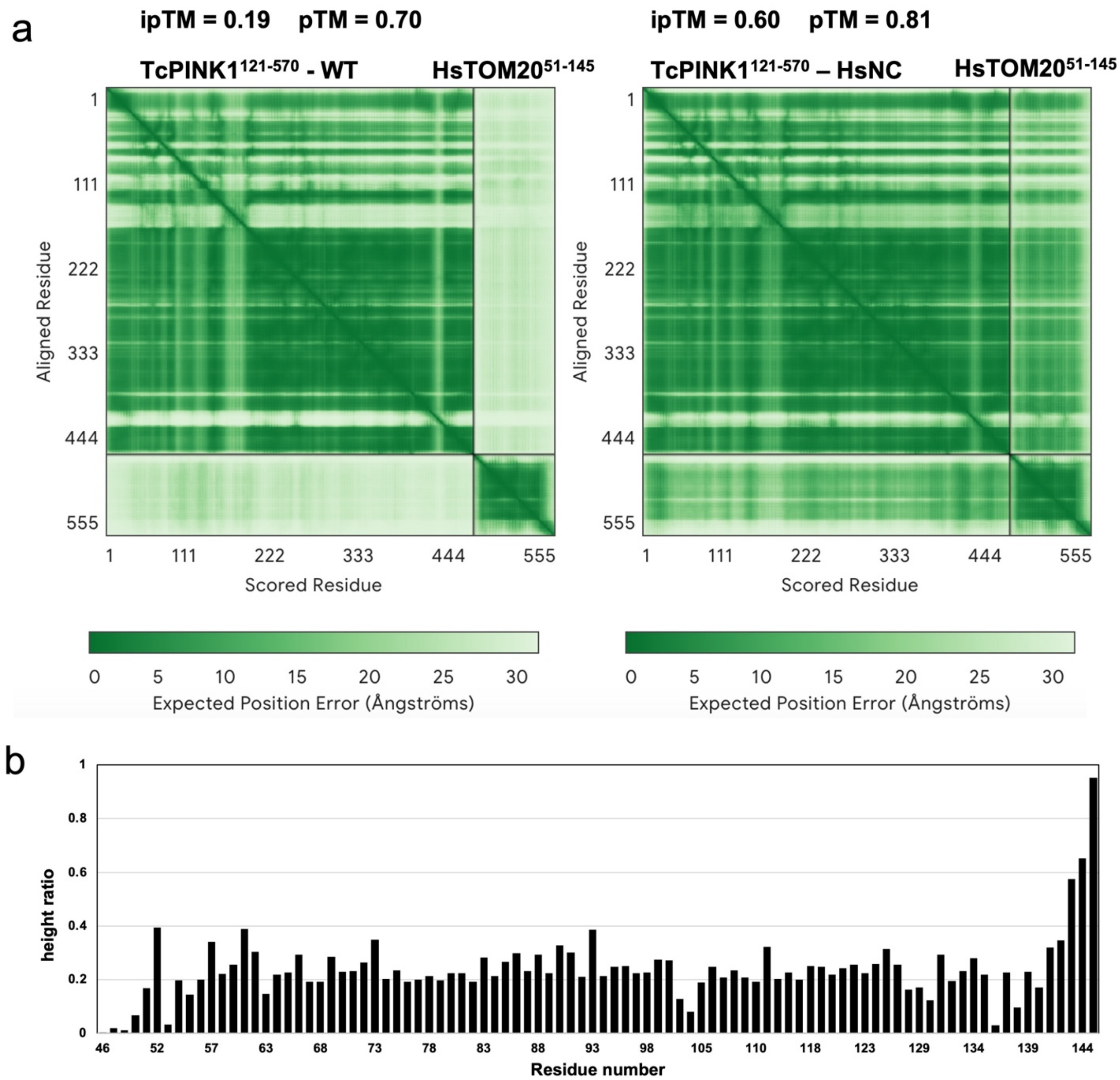


**Figure S3. Humanization of the TcPINK1 NTE–CTE interface promotes interaction with HsTOM20**. **(a)** AlphaFold3 predicted aligned error (PAE) plots for HsTOM20 (51–145) modelled with the humanized $TcPINK1^{HsNC}$ variant (left) or the wild-type TcPINK1 (121–570) (right). **(b)** NMR peak-height ratios for individual HsTOM20 residues following addition of $TcPINK1^{HsNC}$, showing residue-specific line broadening upon complex formation.

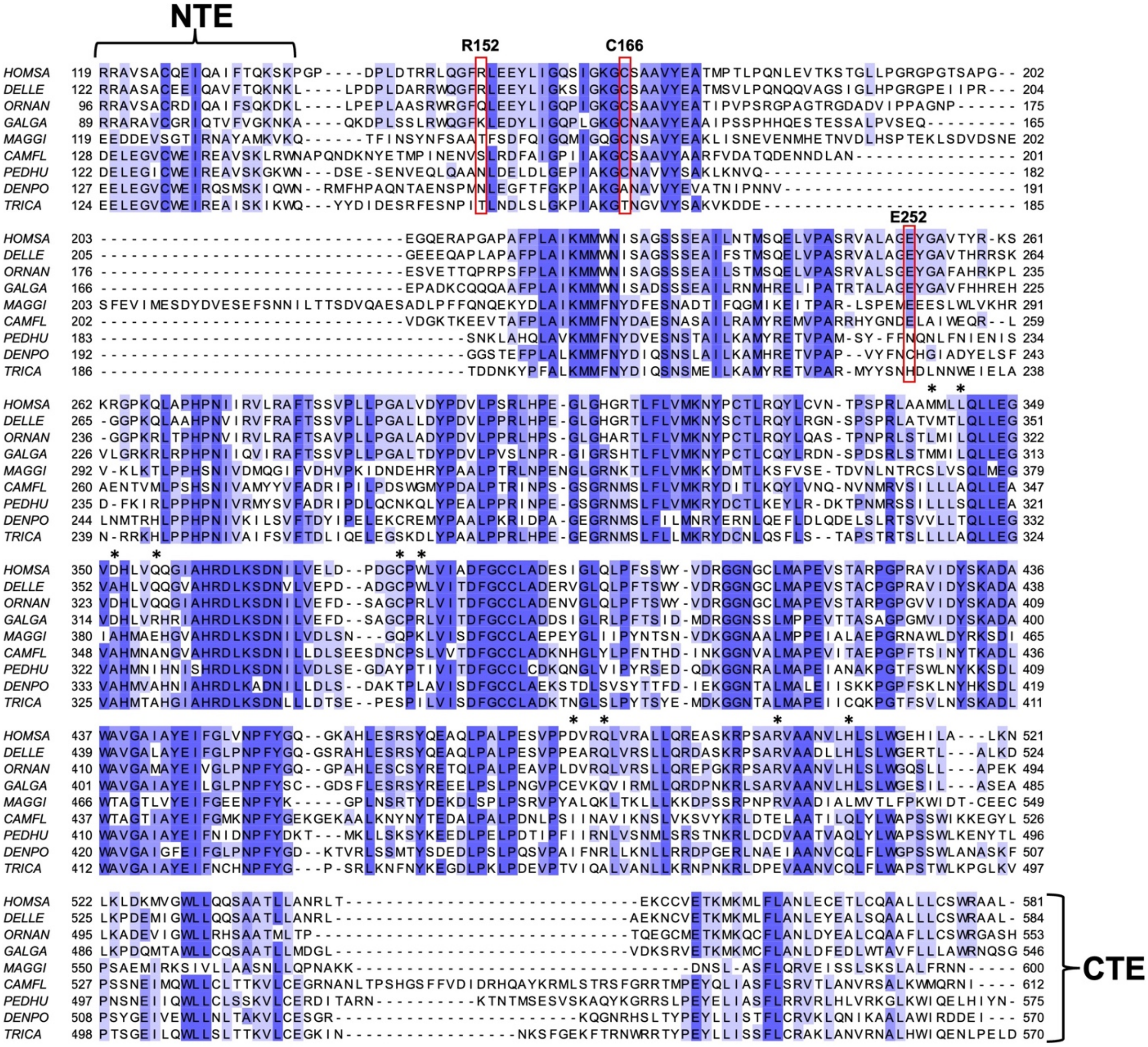


**Figure S4. Sequence conservation of residues examined across PINK1 orthologues**. Multiple sequence alignment of the PINK1 regions containing the NTE–CTE interface and residues investigated experimentally. The positions corresponding to the additional mutations in chimera 1 are annotated with an asterisk. Variants are boxed in red.

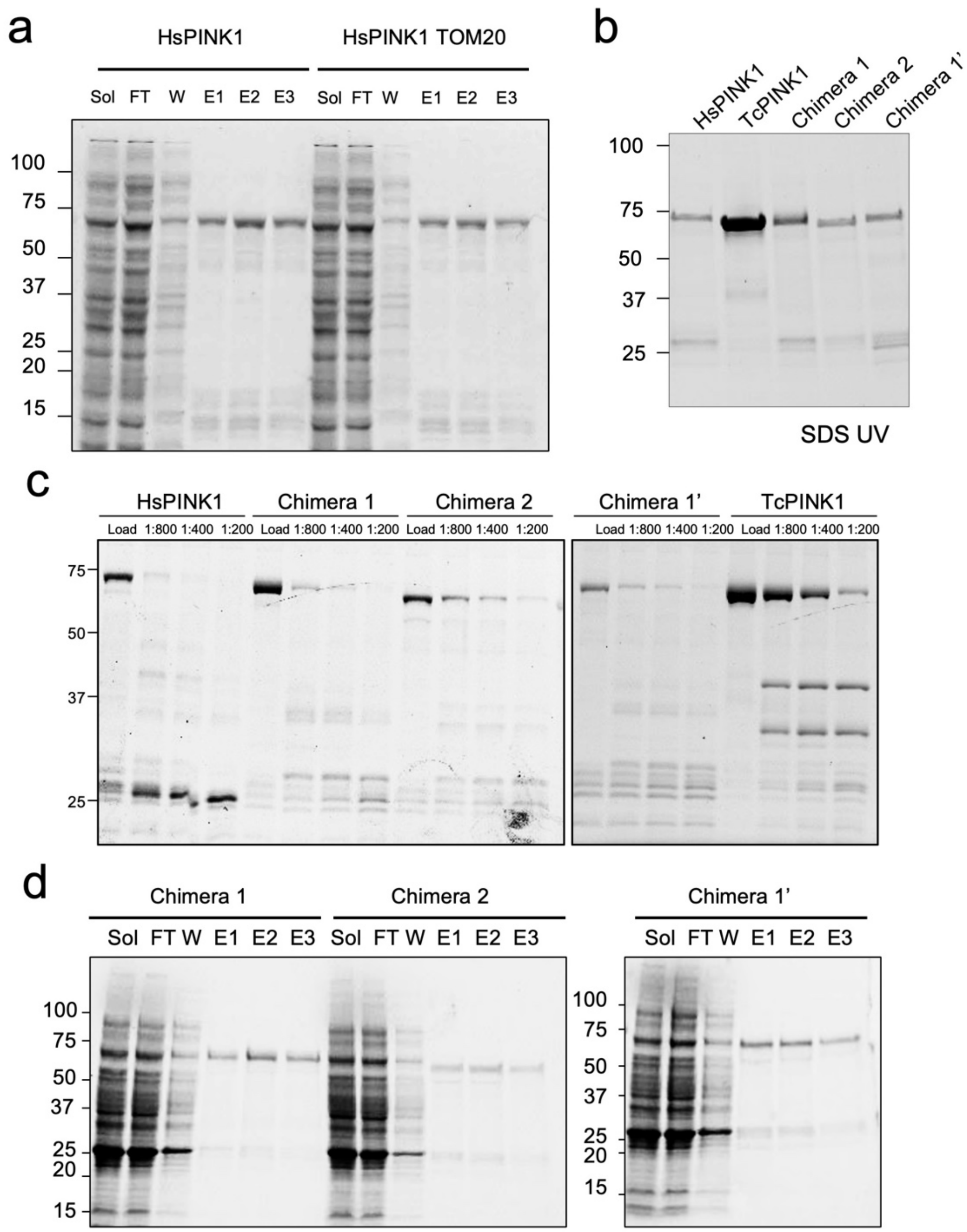


**Figure S5. Recombinant purification and proteolytic stability of HsPINK1 and engineered PINK1 chimeras**. **(a)** GST affinity purification of HsPINK1 expressed alone or co-expressed with HsTOM20. Soluble lysate (Sol), flow-through (FT), wash (W), and sequential glutathione elution fractions (E1–E3) are shown. **(b)** GST affinity purification profiles of HsPINK1, TcPINK1, Chimera 1, Chimera 2, and Chimera 1′ expressed in *E. coli*. Soluble lysate, flow-through, wash, and elution fractions are shown. **(c)** Limited proteolysis of HsPINK1, Chimera 1, Chimera 2, Chimera 1′ and TcPINK1. Purified proteins were incubated with increasing concentrations of elastase, and digestion products were resolved by SDS–PAGE. “Load” indicates untreated protein. **(d)** SDS–PAGE analysis of the purification of Chimera 1, Chimera 2, and Chimera 1′.

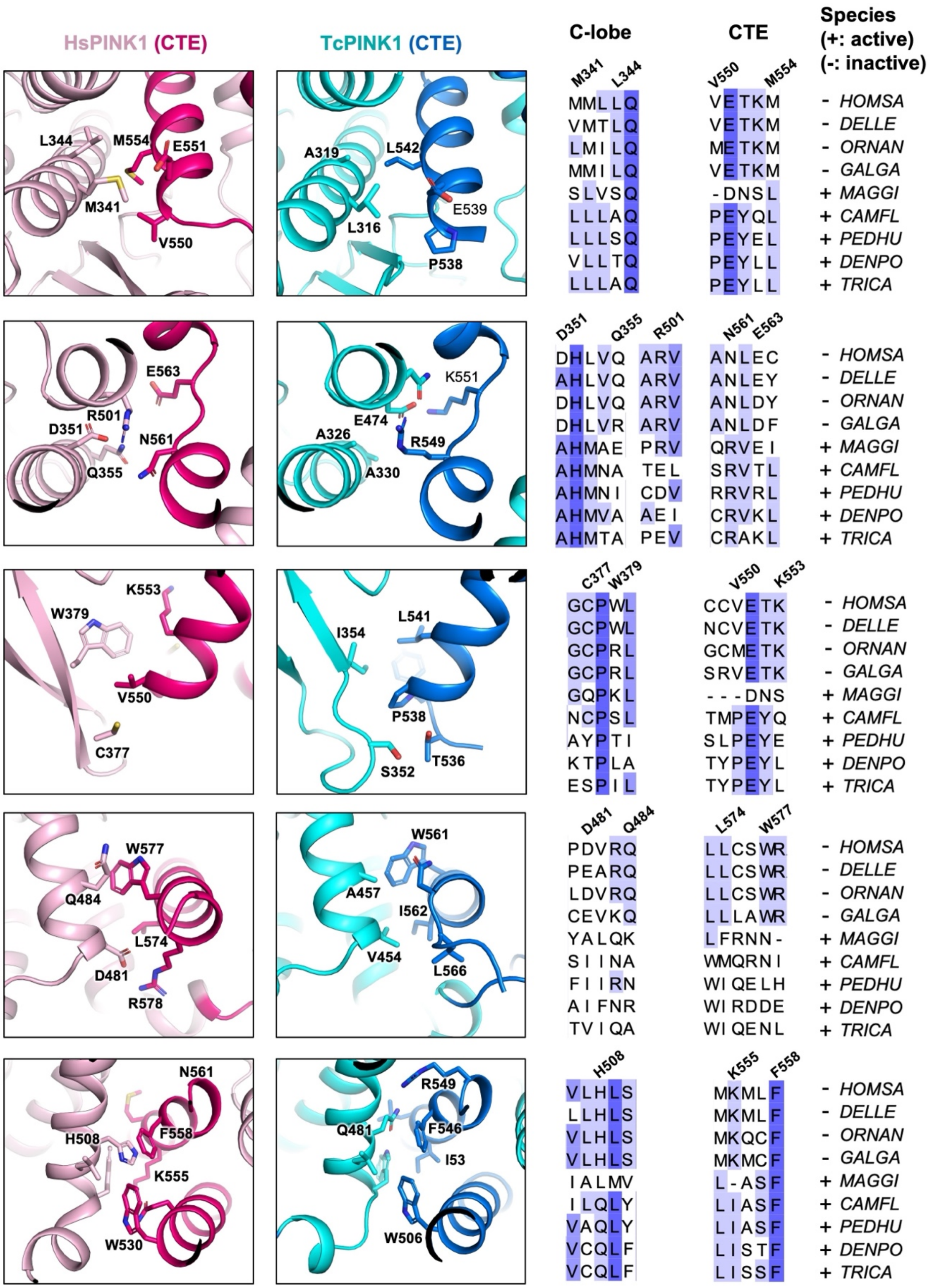


**Figure S6. Structural comparison of the PINK1 CTE–kinase C-lobe interface across vertebrate and invertebrate orthologues**. Predicted interactions between the CTE and kinase C-lobe were analysed across the nine PINK1 orthologues tested experimentally. Representative CTE–C-lobe contacts are shown, with hydrogen-bonding and salt-bridge residues indicated. Experimentally active and inactive recombinant orthologues are denoted by + and −, respectively.

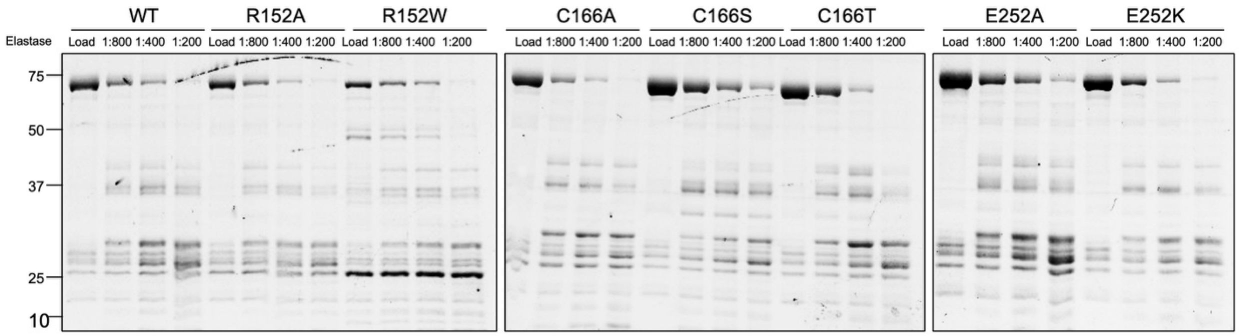


**Figure S7. Proteolytic stability of Chimera 1 variants**. Limited proteolysis of purified WT Chimera 1 and the indicated R152, C166, and E252 variants. Proteins were incubated with increasing concentrations of elastase (enzyme dilutions 1:800, 1:400, and 1:200), and digestion products were resolved by SDS–PAGE. Untreated protein is shown as the loading control.

**Table S1. Sequence identity of TOM20 and PINK1 NTE and CTE regions across vertebrate and invertebrate species.** Percentage sequence identity relative to *Homo sapiens* is shown for TOM20, the PINK1 N-terminal extension (NTE), αK-containing C-terminal extension (CTE–αK), and full C-terminal extension (CTE-full) for the indicated species.

| Species | TOM20 | NTE | CTE-aK | CTE-full |
|---|---|---|---|---|
| Vertebrates | | | | |
| *Homo sapiens* | 100.00% | 100.00% | 100.00% | 100.00% |
| *Acipenser oxyrinchus* | 94.74% | 42.42% | 43.48% | 33.33% |
| *Albula goreensis* | 96.05% | 42.42% | 43.48% | 34.85% |
| *Apteryx owenii* | 69.74% | 10.53% | 73.91% | 66.67% |
| *Betta splendens* | 96.05% | 36.36% | 37.50% | 26.87% |
| *Delphinapterus leucas* | 100% | 83.33% | 82.61% | 86.44% |
| *Eublepharis macularius* | 97.37% | 31.43% | 69.57% | 52.94% |
| *Gallus gallus* | 100% | 24.24% | 60.87% | 56.67% |
| *Lepisosteus oculatus* | 96.05% | 42.42% | 39.13% | 28.99% |
| *Moschus moschiferus* | 100% | 33.33% | 95.65% | 80.95% |
| *Ornithorhynchus anatinus* | 100% | 73.33% | 52.17% | 55.38% |
| *Pavo cristatus* | 78.95% | 24.24% | 60.87% | 56.67% |
| *Podarcis muralis* | 100% | 34.29% | 60.87% | 55.00% |
| Invertebrates | | | | |
| *Acanthosepion pharaonis* | 28.05% | 13.89% | 12.50% | 8.82% |
| *Acrobeloides nanus* | 44.74% | 6.45% | 8.16% | 7.89% |
| *Adineta ricciae* | 44.74% | 5.71% | 12.50% | 18.03% |
| *Araneus ventricosus* | 60.53% | 0.00% | 0.00% | 2.83% |
| *Branchiostoma belcheri* | 78.95% | 27.27% | 31.03% | 27.27% |
| *Caenorhabditis elegans* | 44.74% | 12.90% | 6.25% | 4.49% |
| *Camponotus floridanus* | 65.79% | 16.67% | 9.43% | 10.00% |
| *Dendroctonus ponderosae* | 64.47% | 13.89% | 20.00% | 14.93% |
| *Ditylenchus dipsaci* | 22.37% | 12.90% | 4.08% | 7.76% |
| *Dracunculus medinensis* | 48.68% | 9.68% | 0.00% | 6.12% |
| *Drosophila melanogaster* | 56.58% | 16.67% | 13.51% | 14.86% |
| *Exocentrus adspersus* | 65.79% | 16.67% | 10.00% | 10.34% |
| *Fopius arisanus* | 60.53% | 16.67% | 9.09% | 9.68% |
| *Heterorhabditis bacteriophora* | 55.26% | 9.68% | 8.16% | 5.88% |
| *Holothuria leucospilota* | 68.42% | 7.69% | 11.54% | 18.99% |
| *Larinioides sclopetarius* | 60.53% | 10.81% | 16.13% | 14.71% |
| *Magallana gigas* | 65.79% | 11.43% | 15.38% | 14.29% |
| *Mizuhopecten yessoensis* | 60.53% | 8.33% | 11.11% | 17.19% |
| *Mytilus coruscus* | 65.79% | 5.71% | 8.33% | 11.48% |
| *Oedothorax gibbosus* | 55.26% | 13.51% | 16.13% | 13.24% |
| *Orchesella cincta* | 64.47% | 16.22% | 15.38% | 14.29% |
| *Owenia fusiformis* | 65.79% | 13.89% | 16.22% | 21.62% |
| *Paralvinella palmiformis* | 44.16% | 19.44% | 25.00% | 24.59% |
| *Paramuricea clavata* | 63.16% | 5.00% | 10.00% | 12.86% |
| *Patella caerulea* | 64.47% | 8.82% | 10.71% | 10.77% |
| *Pediculus humanus* | 60.53% | 16.67% | 11.63% | 10.00% |
| *Phallusia mammillata* | 69.74% | 10.26% | 17.39% | 14.29% |
| *Plectus sambesii* | 53.95% | 6.45% | 0.00% | 3.51% |
| *Porites lobata* | 55.26% | 0.00% | 16.67% | 20.90% |
| *Priapulus caudatus* | 56.58% | 7.89% | 6.06% | 7.14% |
| *Saccoglossus kowalevskii* | 73.68% | 10.81% | 12.50% | 11.59% |
| *Scylla olivacea* | 75.00% | 10.81% | 10.87% | 12.05% |
| *Steinernema glaseri* | 53.95% | 9.68% | 4.08% | 8.14% |
| *Strigamia maritima* | 72.37% | 16.22% | 12.90% | 8.82% |
| *Tribolium castaneum* | 60.53% | 13.89% | 11.11% | 10.96% |
